# The Novel Chimeric Multi-Agonist Peptide (GEP44) Reduces Energy Intake and Body Weight in Male and Female Diet-Induced Obese Mice in a Glucagon-Like Peptide-1 Receptor-Dependent Manner

**DOI:** 10.1101/2024.05.17.594690

**Authors:** James E. Blevins, Mackenzie K. Honeycutt, Jared D. Slattery, Matvey Goldberg, June R. Rambousek, Edison Tsui, Andrew D. Dodson, Kyra A. Shelton, Therese S. Salemeh, Clinton T. Elfers, Kylie S. Chichura, Emily F. Ashlaw, Sakeneh Zraika, Robert P. Doyle, Christian L. Roth

## Abstract

We recently reported that a novel chimeric peptide (GEP44) targeting both the glucagon-like peptide-1 receptor (GLP-1R) and neuropeptide Y1- and Y2 receptor (Y1R and Y2R) reduced energy intake and body weight (BW) in diet-induced obese (DIO) rats. We hypothesized that GEP44 reduces energy intake and BW primarily through a GLP-1R dependent mechanism. To test this hypothesis, GLP-1R^+/+^ mice and GLP-1R null (GLP-1R^-/-^) mice were fed a high fat diet for 4 months to elicit diet-induced obesity prior to undergoing a sequential 3-day vehicle period, 3-day drug treatment (5, 10, 20 or 50 nmol/kg; GEP44 vs the selective GLP-1R agonist, exendin-4) and a 3-day washout. Energy intake, BW, core temperature and activity were measured daily. GEP44 (10, 20 and 50 nmol/kg) reduced BW after 3-day treatment in DIO male GLP-1R^+/+^ mice by - 1.5±0.6, -1.3±0.4 and -1.9±0.4 grams, respectively (*P*<0.05), with similar effects being observed in female GLP-1R^+/+^ mice. These effects were absent in male and female DIO GLP-1R^-/-^ mice suggesting that GLP-1R signaling contributes to GEP44-elicited reduction of BW. Further, GEP44 decreased energy intake in both male and female DIO GLP-1R^+/+^ mice, but GEP44 appeared to produce more consistent effects across multiple doses in males. In GLP-1R^-/-^ mice, the effects of GEP44 on energy intake were only observed in males and not females, suggesting that GEP44 may reduce energy intake, in part, through a GLP-1R independent mechanism in males. In addition, GEP44 reduced core temperature and activity in both male and female GLP-1R^+/+^ mice suggesting that it may also reduce energy expenditure. Lastly, we show that GEP44 reduced fasting blood glucose in DIO male and female mice through GLP-1R. Together, these findings support the hypothesis that the chimeric peptide, GEP44, reduces energy intake, BW, core temperature, and glucose levels in male and female DIO mice primarily through a GLP-1R dependent mechanism.

## Introduction

Obesity is a major worldwide health concern as it increases the risk of cardiovascular disease, obstructive sleep apnea, cancer, osteoarthritis, depression, COVID-19 related hospitalizations and type 2 diabetes. According to the NCD Risk Factor Collaboration, more than one billion people are obese worldwide [1]. Approximately 1 in 2 US adults are predicted to have obesity by 2030 [2] and the costs to treat obesity in the US are estimated to be approximately 3 trillion/year by 2030 [3]. Current pharmacotherapies to treat obesity are either not well tolerated or are associated with unwanted and/or adverse side effects (i.e. nausea, vomiting, diarrhea, depression, and sleep disturbance) that limit their long-term use, highlighting the need for more effective treatment options. While the more recently developed monotherapies to treat obesity such as the long-acting glucagon-like peptide-1 receptor (GLP-1R) agonists, liraglutide and semaglutide [4], produce pronounced effects on weight loss relative to previous analogues, they can be associated with gastrointestinal (GI) side effects.

Recent studies suggest that combination therapy aimed at suppressing energy intake and/or increasing energy expenditure at low-dose or subthreshold dose combinations may be more effective for producing sustained weight loss than monotherapy [5] and may minimize the potential for unwanted side effects. The overall ineffectiveness of monotherapies to evoke prolonged weight loss in humans with obesity is assumed to occur, in part, by recruitment of robust orexigenic mechanisms that drive energy intake and decrease energy expenditure, resulting in body weight (BW) gain and thus preventing further weight loss. Two of the currently available US Food and Drug Administration (FDA)-approved weight loss drugs include combination therapies, namely Qsymia (phentermine + topiramate) and Contrave (bupoprion + naltrexone) [6], which have resulted in 10.9% [7] and 6.4% [8] weight loss, respectively.

In addition to combination therapy, a recent innovative approach involves the targeting of two or more signaling pathways using a single compound such as monomeric multi-agonists (dual- or triple-agonists) based on glucose-dependent insulinotropic polypeptide (GIP) and GLP-1R, with and without glucagon receptor (GCGR) agonism. One such drug, tirzepatide (Zepbound^TM^), which targets both GLP-1R and GIP, was found to elicit a robust 20.9% and 25.3% weight loss in humans with obesity over 72- [9] and 88-week periods [10], and was recently approved by the FDA for weight management. Recent data indicate that the triple-agonist, retatrutide, which targets GLP-1R, GIP and glucagon receptors, was able to elicit 24.2% weight loss over 48-week period [11] (2 to 12 mg). While such therapies show considerable promise for effective and sustained reduction of BW during drug treatment, there are still adverse gastrointestinal side effects including nausea, diarrhea, abdominal pain and vomiting [9], leading, in some cases, to the discontinuation of the drug in up to 7.1% of participants [9]. Thus, there remains an urgent need to develop newer and equally effective anti-obesity treatment strategies that eliminate the adverse side effects particularly for patients that may need long-term treatment.

We recently designed the novel chimeric peptide, GEP44, which binds to GLP-1R, Y1R and Y2R [12]. We found that systemic administration of GEP44 reduced energy intake and BW in both lean [13] and diet-induced obese rats [12; 13]. Importantly, GEP44 also reduced energy intake at doses that were not associated with significant pica behavior (kaolin intake) in rats [13] or emesis in musk shrews [13]. A critical remaining question is whether GEP44 reduces energy intake, BW, impacts thermogenesis and improves glucose homeostasis in a GLP-1R dependent manner, either alone or in-part. Given the role of GLP-1R in the control of energy intake and BW, we hypothesized that chimeric peptide GEP44 reduces energy intake and BW through a GLP-1R dependent mechanism. To test this hypothesis, we determined the extent to which GEP44 reduces energy intake and BW, impacts core temperature (core temperature as surrogate marker of energy expenditure) and gross motor activity and improves fasting blood glucose in male and female DIO mice that lack GLP-1R (GLP- 1R^-/-^) relative to age-matched cohorts of male and female GLP-1R^+/+^ mice. Male and female mice were also weight-matched within a given cohort of GLP-1R^-/-^ and GLP- 1R^+/+^ mice prior to treatment. The selective GLP-1R agonist, exendin-4, was included as a control to assess GLP-1R mediated effects on energy intake [14].

## Methods

### Animals

GLP-1R^+/-^ mice were initially obtained from Dr. Daniel Drucker (University of Toronto, Canada) and bred by Dr. Sakeneh Zraika at the Veterans Affairs Puget Sound Health Care System (VAPSHCS) to obtain GLP-1R^+/+^ and GLP-1R^-/-^ (GLP-1R null) mice. Adult male and female mice (age: ∼5.5-10 weeks) weighed, on average, 18.04±0.04 grams and were 10.9±0.03% fat at the time of body composition measurements prior to diet intervention. Mice were initially maintained on a chow diet [PicoLab^®^ Rodent Diet 20 (5053)] (LabDiet^®^, St. Louis, MO; 13% kcal from fat). Mice were subsequently placed on a high fat diet (HFD) [60% kcal from fat; Research Diets, Inc., D12492i, New Brunswick, NJ] for approximately 4 months at which time body composition measurements were completed (**Table 1**). Mice weighed, on average, 39.3±0.04 grams and were 39.8±0.03% fat at the time of body composition measurements prior to drug intervention. All animals were housed individually in Plexiglas cages in a temperature-controlled room (22±2°C) under a 12:12-h light-dark cycle. All mice were maintained on a 11 a.m./11 p.m. reverse light cycle (lights off at 11 a.m./lights on at 11 p.m.). Mice had *ad libitum* access to water and HFD. The research protocols were approved both by the Institutional Animal Care and Use Committee of the Veterans Affairs Puget Sound Health Care System (VAPSHCS) and the University of Washington in accordance with NIH Guidelines for the Care and Use of Animals.

**Table 1.**
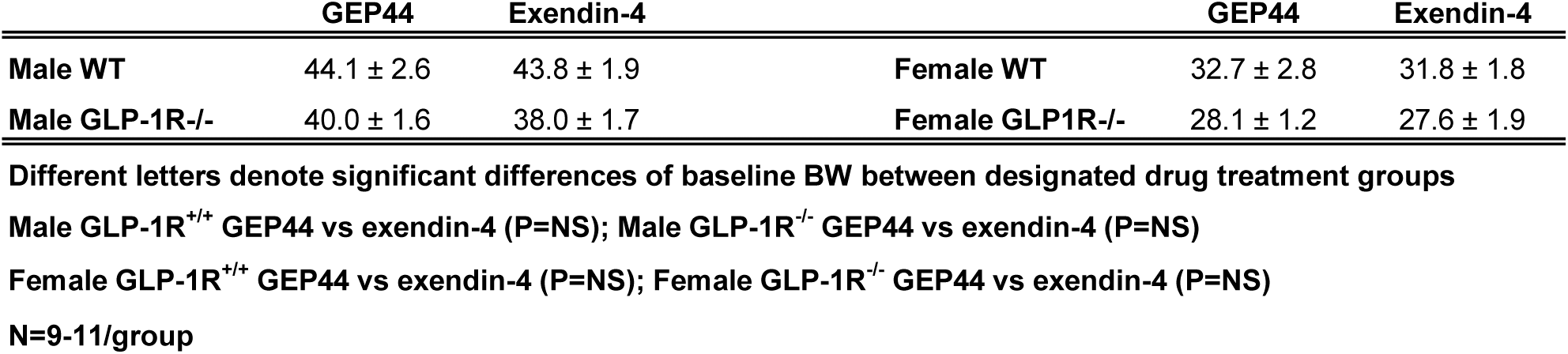
Body weight of Male and Female GLP-1R^+/+^ and GLP-1R^-/-^ DIO mice prior to treatment intervention at study onset (N=9-11/group). Data are expressed as mean ± SEM.

### Drug Preparation

GEP44 was synthesized in the Doyle lab as previously described [12]. Fresh solutions of GEP44 and exendin-4 (ENZO; Farmingdale, NY) were prepared, frozen and thawed prior to the onset of each experiment (**Study 1**).

### Implantation of G2 E-Mitter telemetry devices into abdominal cavity

At the age of 27-31.5 weeks (average age 28.8±0.01 weeks), animals were anesthetized with isoflurane and subsequently underwent a sham surgery (no implantation) or received implantations of a sterile G2 E-Mitter (15.5 mm long x 6.5 mm wide; Starr Life Sciences Company) into the intraperitoneal cavity. The abdominal opening was closed with 5-0 Vicryl® absorbable suture and the skin was closed with Nylon sutures (5-0). Vetbond glue was used to seal the wound and bind any tissue together between the sutures. Sutures were removed within two weeks after the G2 E- Mitter implantation. All G2 E-Mitters were confirmed to have remained within the abdominal cavity at the conclusion of the study.

### Acute SC injections of GEP44 and Exendin-4

GEP44 (or saline vehicle; 3 mL/kg injection volume) or exendin-4 were administered immediately prior to the start of the dark cycle following 2 hours of food deprivation. Mice underwent all treatments (unless otherwise noted). The study design consisted of sequential rounds of a 3-day baseline phase (vehicle treated), a 3-day treatment phase (single dose repeated over 3 days), and a washout phase (3 days). The 3-day treatment phase consisted of a dose escalation design beginning with the low dose (5 nmol/kg) over week 1 and ending with the high dose (50 nmol/kg) over the final week of the study (5, 10, 20 and 50 nmol/kg; GEP44 vs the selective GLP-1R agonist, exendin-4). A separate set of age- and weight-matched mice were treated with vehicle in place of drug in order to determine the impact of drug treatment on tail vein glucose, plasma hormones and thermogenic gene expression. Body weight was assessed daily approximately 3-h prior to the start of the dark cycle.

### Body Composition

Determinations of lean body mass and fat mass were made on conscious mice by quantitative magnetic resonance using an EchoMRI 4-in-1-700^TM^ instrument (Echo Medical Systems, Houston, TX) at the VAPSHCS Rodent Metabolic Phenotyping Core.

## Study Protocols

### Changes of BW and Energy Intake

DIO GLP-1R^+/+^ and GLP-1R^-/-^ mice underwent a 3- day vehicle period, 3-day dose escalation drug treatment (5, 10, 20 and 50 nmol/kg; GEP44 vs the selective GLP-1R agonist, exendin-4) and a 3-day washout in sequential order. The selective GLP-1R agonist, exendin-4, was included as a control to assess GLP-1R mediated effects on energy intake [14]. BW and energy intake were measured daily. Energy intake data was averaged throughout each sequential 3-day vehicle and 3-day drug treatment period.

### Changes of Core Temperature and Gross Motor Activity

Telemetry recordings of core temperature (surrogate marker of energy expenditure) and gross motor activity were measured from each mouse in the home cage immediately prior to injections and for a 6-, 12-, and 24-h period after injections. Core temperature and gross motor activity were recorded every 15 sec. The last hour of the light cycle (during which time energy intake, BW and drug administration occurred) was excluded from the telemetry analysis.

### Tissue Collection for Quantitative Real-Time PCR (qPCR)

Tissue [interscapular brown adipose tissue (IBAT) was collected from 3-h fasted mice at 2-h post-injection. Mice were euthanized with an overdose of ketamine cocktail at 2-h post-injection. Tissue was rapidly removed, wrapped in foil and frozen in liquid N2. Samples were stored frozen at -80°C until analysis.

#### Blood collection

Blood samples [up to 1 mL] were collected by cardiac stick in chilled K2 EDTA Microtainer Tubes (Becton-Dickinson, Franklin Lakes, NJ). Whole blood was centrifuged at 6,000 rpm for 1.5-min at 4°C; plasma was removed, aliquoted and stored at −80°C for subsequent analysis.

#### Blood Glucose Measurements

Blood was collected in a subset of mice for glucose measurements by tail vein nick and measured using a glucometer (AlphaTRAK 2, Abbott Laboratories, Abbott Park, IL) [15; 16].

#### Plasma hormone measurements

Plasma leptin and insulin were measured using electrochemiluminescence detection [Meso Scale Discovery (MSD^®^), Rockville, MD] using established procedures [16; 17]. Intra-assay coefficient of variation (CV) for leptin was 5.9% and 1.6% for insulin (pending assay data). The range of detectability for the leptin assay is 0.137-100 ng/mL and 0.069-50 ng/mL for insulin (pending assay data). The data were normalized to historical values using a pooled plasma quality control sample that was assayed in each plate.

#### Tissue collection for quantitative real-time PCR (qPCR)

Tissue (IBAT) was collected from a subset of 3-h fasted mice (at 2-h post-injection). Mice were euthanized with an overdose of ketamine cocktail prior to tissue collection. IBAT was collected within a 5-h window towards the end of the light cycle (9:00 a.m.- 2:00 p.m.) as previously described in DIO CD^®^ IGS/Long-Evans rats and C57BL/6J mice [15; 16; 18]. Tissue was rapidly removed, wrapped in foil and frozen in liquid nitrogen. Samples were stored frozen at -80°C until analysis.

#### qPCR

RNA extracted from samples of IBAT was analyzed using the RNeasy Lipid Mini Kit (Qiagen Sciences Inc, Germantown, MD) followed by reverse transcription into cDNA using a high-capacity cDNA archive kit (Applied Biosystems, Foster City, CA).

Quantitative analysis for relative levels of mRNA in the RNA extracts was measured in duplicate by qPCR on an Applied Biosystems 7500 Real-Time PCR system (Thermo Fisher Scientific, Waltham, MA) using the following TaqMan® probes (Thermo Fisher Scientific Gene Expression Assay probes): mouse Nono (catalog no.

Mm00834875_g1), mouse UCP-1 (catalog no. Mm01244861_m1), mouse type 2 deiodinase (D2) (Dio2; catalog no. Mm00515664_m1), mouse G-protein coupled receptor 120 (Gpr120; catalog no. Mm00725193_m1), mouse cell death-inducing DNA fragmentation factor alpha-like effector A (Cidea; catalog no. Mm00432554_m1) and mouse peroxisome proliferator-activated receptor gamma coactivator 1 alpha (Ppargc1a; catalog no. Mm01208835_m1). Relative amounts of target mRNA were determined using the Comparative CT or 2-^ΔΔC^T method [19] following adjustment for the housekeeping gene, Nono. Specific mRNA levels of all genes of interest were normalized to the cycle threshold value of Nono mRNA in each sample and expressed as changes normalized to controls (vehicle treatment).

## Statistical Analyses

All results are expressed as mean ± SE. Comparisons between multiple groups involving between subject design were made using one-way ANOVA, followed by a post-hoc Fisher’s least significant difference test. Comparisons involving within-subjects designs were made using a one-way repeated-measures ANOVA followed by a post-hoc Fisher’s least significant difference test. Analyses were performed using the statistical program SYSTAT (Systat Software, Point Richmond, CA). Differences were considered significant at *P*<0.05, 2-tailed.

## Results

The overall goal of these studies was to use the GLP-1R^-/-^ mouse as a strategy to determine the extent to which chimeric peptide, GEP44, reduces BW and energy intake, impacts core temperature (as surrogate for energy expenditure) and activity, and improves glucose homeostasis through the GLP-1R in both male and female DIO mice. In addition, we incorporated use of the selective GLP-1R agonist, exendin-4, to assess GLP-1R mediated effects on BW, energy intake, core temperature, activity, and glucose levels in male and female DIO mice.

### Baseline BW-matching in Male and Female DIO GLP-1R^+/+^ and GLP-1R^-/-^ Mice at Study Onset (post-dietary intervention/pre-drug treatment)

By design, there were no differences in baseline BW between designated drug treatment groups (GEP44 vs exendin-4) of male GLP-1R^+/+^ mice [F(1,19) = 0.009, P=NS] or male GLP-1R^-/-^ mice [F(1,18) = 0.682, P=NS] prior to drug treatment (**Table 1**). Similarly, there were also no differences in baseline BW between designated treatment groups (GEP44 vs exendin- 4) of female GLP-1R^+/+^ mice [F(1,18) = 0.071, P=NS] or GLP-1R^-/-^ mice [F(1,16) = 0.037, P=NS] prior to drug treatment (**Table 1**).

### Changes in BW and Energy Intake

The goal of these studies was to use the GLP-1^-/-^ mouse as a strategy to determine the extent to which chimeric peptide, GEP44, reduces BW and energy intake through the GLP-1R in both male and female DIO mice. In addition, we incorporated use of the selective GLP-1R agonist, exendin-4, to assess GLP-1R mediated effects on BW and energy intake in male and female DIO mice.

## Body Weight

Both male and female GLP-1R^+/+^ and GLP-1R^-/-^ mice were weight-matched within respective groups prior to treatment onset (**Table 1**). GEP44 treatment reduced cumulative 3-day BW in both male (**Fig. 1A-1B**) and female DIO GLP-1R^+/+^ mice (**Fig. 1C-1D**), but these effects were absent in male and female DIO GLP-1R^-/-^ mice. Specifically, GEP44 (10, 20 and 50 nmol/kg), reduced 3-day BW in DIO male GLP-1R^+/+^ mice by -1.5±0.6, -1.3±0.4 and -1.9±0.4 grams, respectively (**Fig. 1A-1B**; *P*<0.05). GEP44 also reduced 3-day BW (10, 20 and 50 nmol/kg) in female GLP-1R^+/+^ mice by - 0.5±0.5 (*P*=0.053), -1.9±0.4 and -1.3±0.3 grams, respectively (**Fig. 1C-1D**; *P*<0.05).

**Figure 1A-D.**
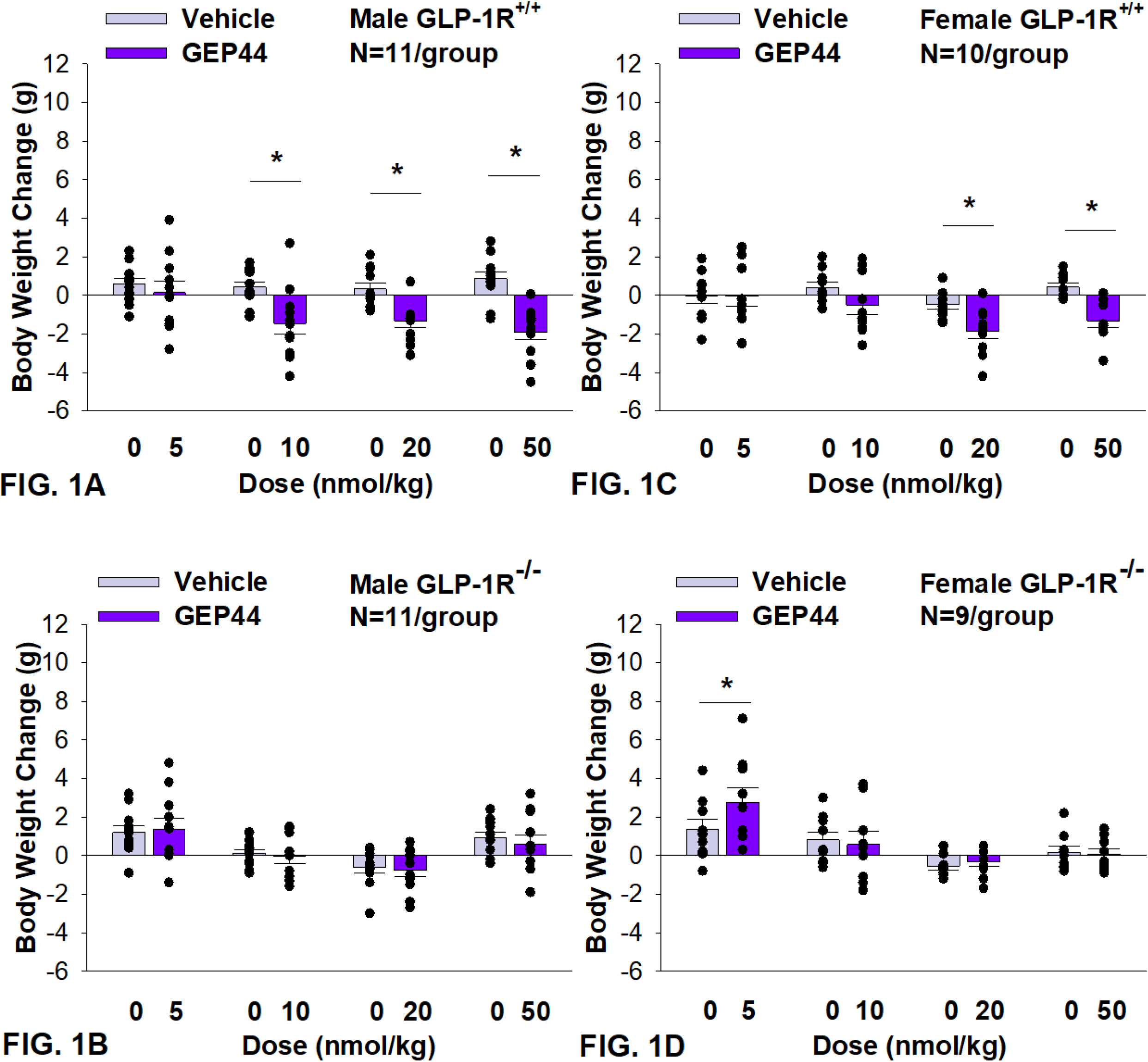
Effects of the chimeric peptide, GEP44, on BW in male and female DIO GLP-1R^+/+^ and GLP-1R^-/-^ mice. Mice were maintained on HFD (60% kcal from fat; N=10-11/group) for approximately 4 months prior to receiving SC injections of vehicle (sterile saline/water) followed by escalating doses of GEP44 (5, 10, 20 and 50 nmol/kg; 3 mL/kg injection volume). *A*, Effect of GEP44 on change in BW in male HFD-fed DIO GLP-1R^+/+^ mice; *B*, Effect of GEP44 on change in BW in male HFD-fed DIO GLP-1R ^-/-^ mice; *C*, Effect of GEP44 on change in BW in female HFD-fed DIO GLP-1R^+/+^ mice; *D*, Effect of GEP44 on change in BW in female HFD-fed DIO GLP-1R^-/-^ mice. Data are expressed as mean ± SEM. \**P*<0.05 GEP44 vs. vehicle.

Similarly, the selective GLP-1R agonist, exendin-4, reduced BW only in male (**Fig. 2A-2B;** *P*<0.05) and female DIO GLP-1R^+/+^ mice (**Fig. 2C-2D**; *P*<.05) but not in male and female DIO GLP-1R ^-/-^ mice. Specifically, exendin-4 (5, 10, 20 and 50 nmol/kg) reduced BW in DIO male GLP-1R^+/+^ mice by -0.7±0.5, -1.0±0.4, -1.4±0.3, and -1.5±0.2 grams, respectively (**Fig. 1A-1B**; *P*<0.05). Exendin-4 also reduced BW (10, 20 and 50 nmol/kg) in female GLP-1R^+/+^ mice by -0.9±0.6, -0.9±0.3 and -1.5±0.2 grams, respectively (**Fig. 1C-1D**; *P*<0.05).

**Figure 2A-D:**
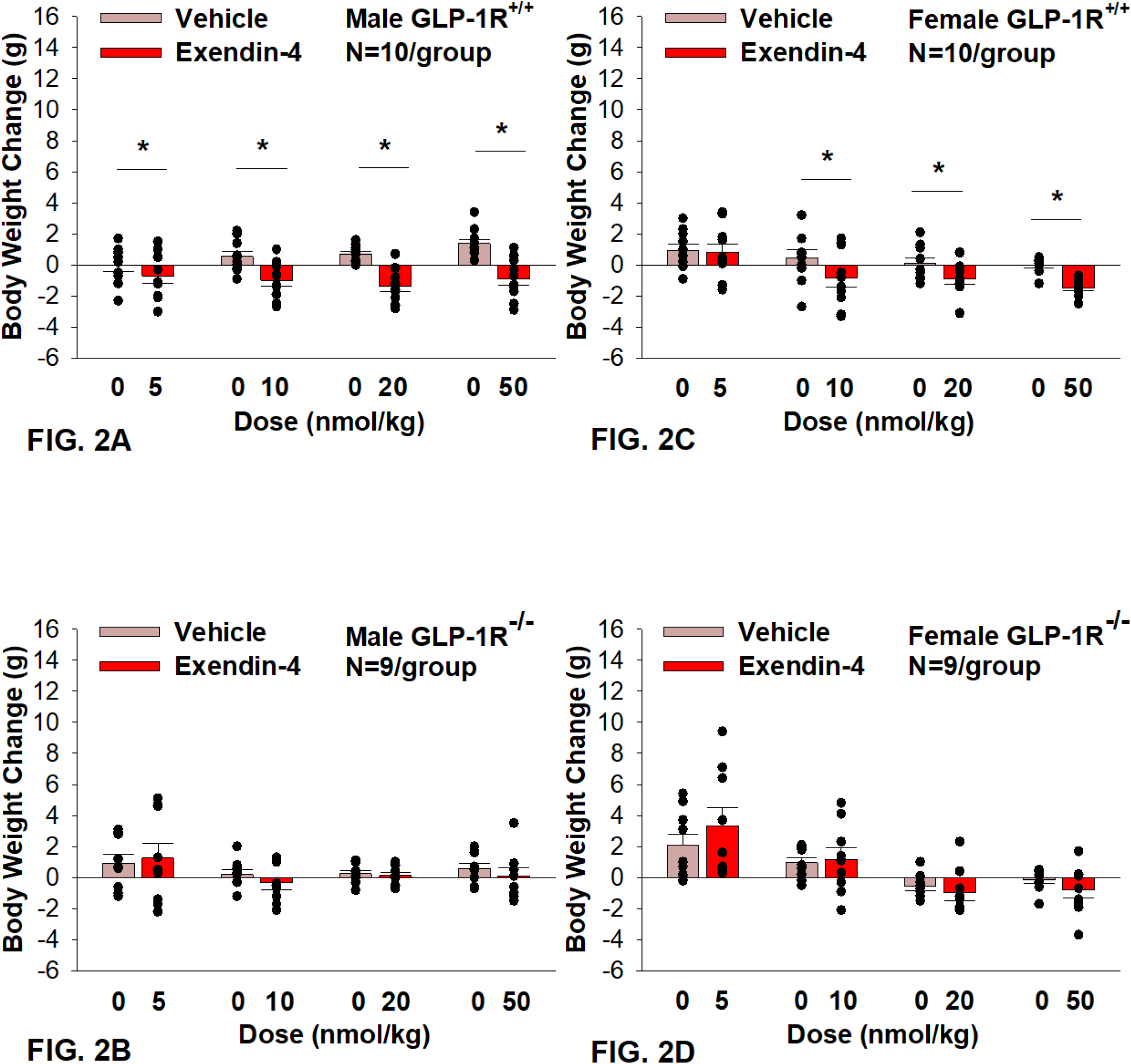
Effects of the selective GLP-1R agonist, exendin-4, on BW in male and female DIO GLP-1R^+/+^ and GLP-1R^-/-^ mice. Mice were maintained on HFD (60% kcal from fat; N=9-11/group) for approximately 4 months prior to receiving SC injections of vehicle (sterile saline/water) followed by escalating doses of exendin-4 (5, 10, 20 and 50 nmol/kg; 3 mL/kg injection volume). *A*, Effect of exendin-4 on change in BW in male HFD-fed DIO GLP-1R^+/+^ mice; *B*, Effect of exendin-4 on change in BW in male HFD-fed DIO GLP-1R^-/-^ mice; *C*, Effect of exendin-4 on change in BW in female HFD-fed DIO GLP-1R^+/+^ mice; *D*, Effect of exendin-4 on change in BW in female HFD-fed DIO GLP- 1R^-/-^ mice. Data are expressed as mean ± SEM. \**P*<0.05 exendin-4 vs. vehicle.

## Energy intake

GEP44 treatment significantly reduced energy intake only in male (**Fig. 3A-3B**) and female DIO GLP-1R^+/+^ mice (**Fig. 3C-3D**) but not in male (with exception of the lowest and highest dose) and female DIO GLP-1R ^-/-^ mice. GEP44 treatment decreased energy intake (10, 20 and 50 nmol/kg); *P*<0.005) in both male and female mice. Similar results on FI were obtained when normalizing to BW (**Supplemental Fig. 1A-D**).

**Figure 3A-D:**
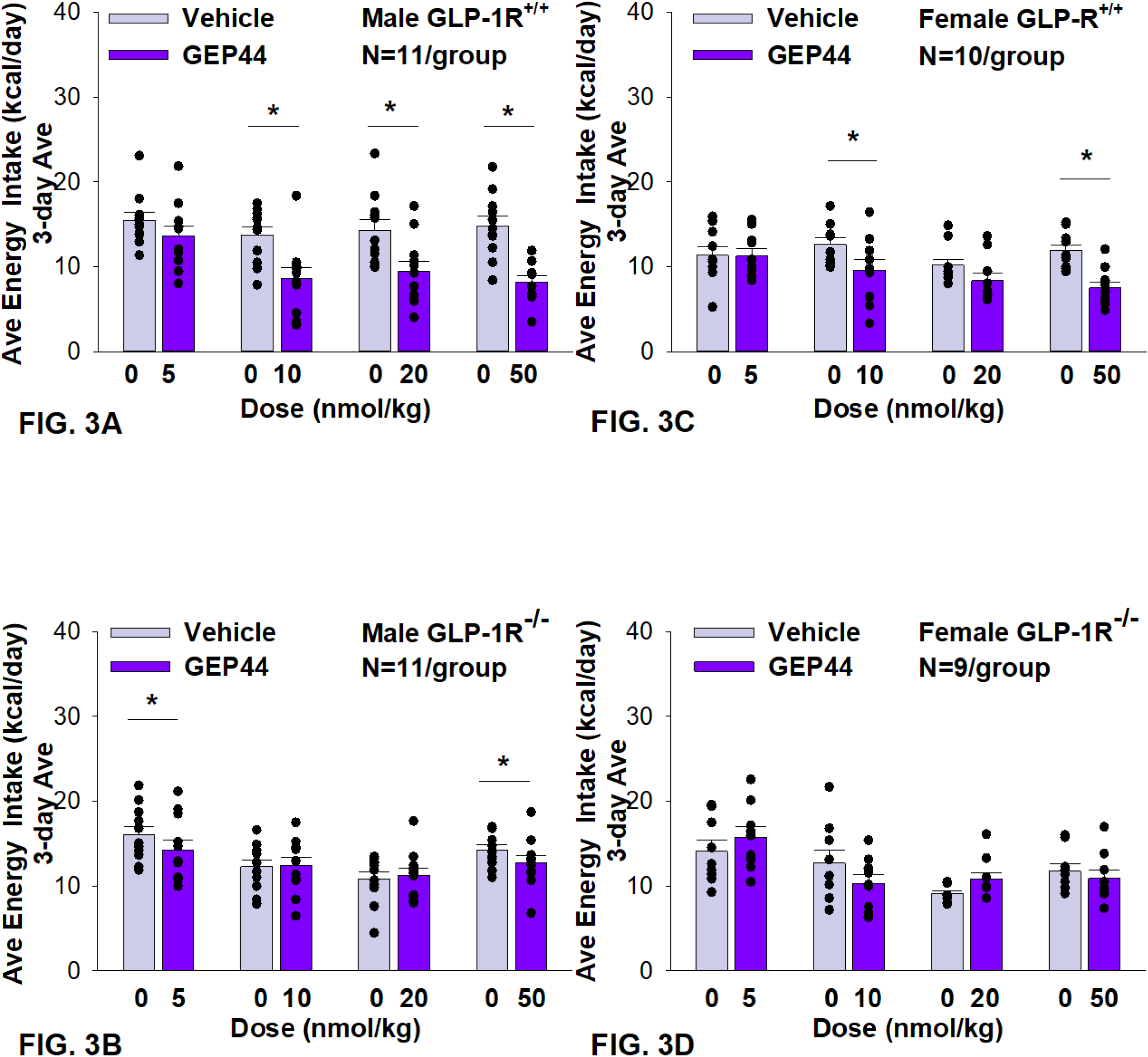
Effects of the chimeric peptide, GEP44, on energy intake (kcal/day) in male and female DIO GLP-1R^+/+^ and GLP-1R ^-/-^ mice. Mice were maintained on HFD (60% kcal from fat; N=10-11/group) for approximately 4 months prior to receiving SC injections of vehicle (sterile saline/water) followed by escalating doses of GEP44 (5, 10, 20 and 50 nmol/kg; 3 mL/kg injection volume). *A*, Effect of GEP44 on energy intake in male HFD-fed DIO GLP-1R^+/+^ mice; *B*, Effect of GEP44 on change on energy intake in male HFD-fed DIO GLP-1R^-/-^ mice; *C*, Effect of GEP44 on energy intake in male HFD-fed DIO GLP-1R^+/+^ mice; *D*, Effect of GEP44 on energy intake in male HFD-fed DIO GLP-1R^-/-^ mice. Data are expressed as mean ± SEM. \**P*<0.05 GEP44 vs. vehicle.

Exendin-4 treatment also significantly reduced energy intake in male (**Fig. 4A-4B**) and female DIO GLP-1R^+/+^ mice (**Fig. 4C-4D**) but not in male (with exception of the lowest dose) and female DIO GLP-1R ^-/-^ mice. Specifically, exendin-4 treatment decreased energy intake (10, 20 and 50 nmol/kg; *P*<0.05) in male GLP-1R^+/+^ mice and also tended to decrease energy intake at the low dose (5 nmol/kg) in male GLP-1R ^-/-^ mice (*P*=0.055). However, exendin-4 reduced energy intake in female GLP-1R^+/+^ mice across all doses (5, 10, 20 and 50 nmol/kg; *P*<0.05).

**Figure 4A-D:**
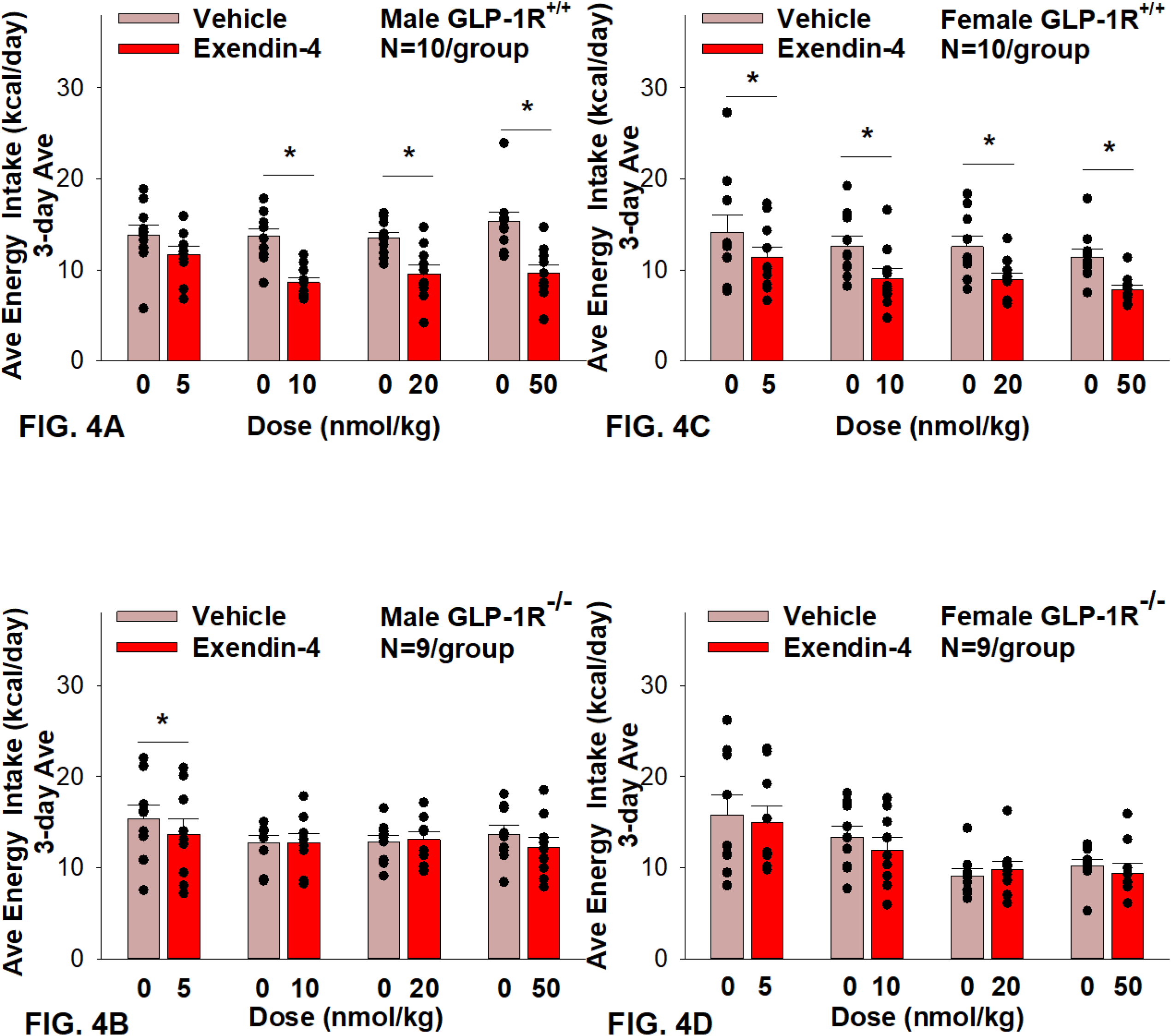
Effects of the selective GLP-1R agonist, exendin-4, on energy intake (kcal/day) in male and female DIO GLP-1R^+/+^ and GLP-1^-/-^ mice. Mice were maintained on HFD (60% kcal from fat; N=9-11/group) for approximately 4 months prior to receiving SC injections of vehicle followed by escalating doses of GEP44 (5, 10, 20 and 50 nmol/kg; 3 mL/kg injection volume). *A*, Effect of GEP44 on energy intake in male HFD-fed DIO GLP-1R^+/+^ mice; *B*, Effect of GEP44 on change on energy intake in male HFD-fed DIO GLP-1R^-/-^ mice; *C*, Effect of GEP44 on energy intake in male HFD-fed DIO GLP-1R^+/+^ mice; *D*, Effect of GEP44 on energy intake in male HFD-fed DIO GLP- 1R^-/-^ mice. Data are expressed as mean ± SEM. \**P*<0.05 exendin-4 vs. vehicle.

Similar results on energy intake were obtained when normalizing to BW (**Supplemental Fig. 2A-D**).

### Changes of Core Temperature and Gross Motor Activity

#### Core temperature

GEP44 and exendin-4 both reduced core temperature over the 6-h post-injection period in male (**Fig. 5A**; *P*<0.05) and female DIO GLP-1R^+/+^ mice (**Fig. 5C**; *P*<0.05). Notably, similar results were observed during the 12-h dark cycle in response to GEP44 in male DIO GLP-1R^+/+^ mice (*P*<0.05; data not shown). However, the effect of GEP44 to reduce core temperature was observed only at the high dose in male GLP-1R^-/-^ mice (50 nmol/kg; **Fig. 5B**; *P*<0.05). GEP44 was unable to reduce core temperature in female GLP-1R^-/-^ mice (**Fig. 5D**; *P*=NS). However, GEP44 produced an unexpected stimulation of core temperature at the low dose in female GLP-1R^-/-^ mice (**Fig. 5D**; *P*<0.05). There was an effect of exendin-4 to reduce core temperature in male (**Fig. 6A**; *P*<0.05) or female GLP-1R^+/+^ mice (**Fig. 6C**; *P*<0.05) but not in GLP-1R^-/-^ mice (**Fig. 6B**; **Fig. 6D**; *P*=NS).

**Figure 5A-D.**
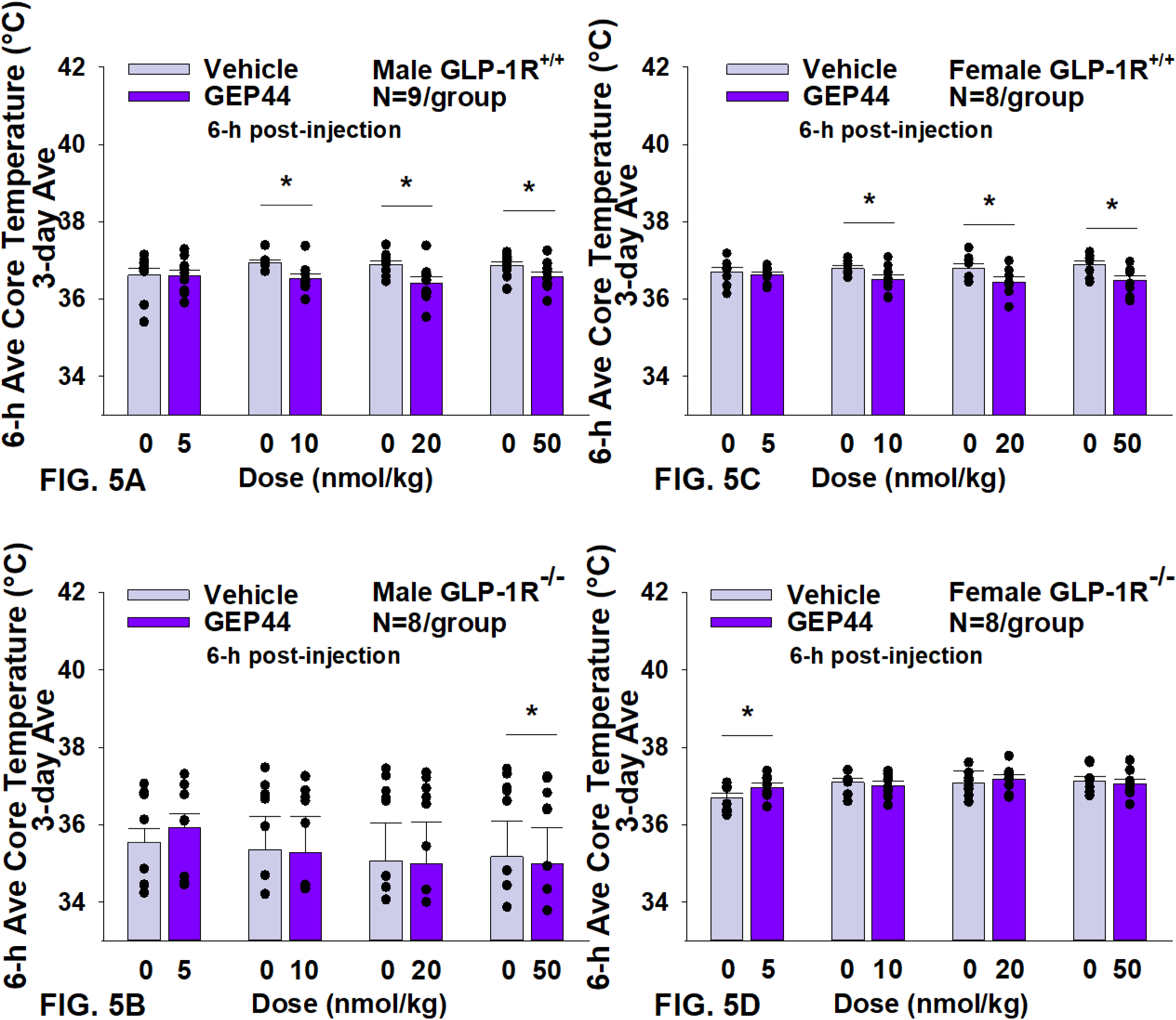
Effects of the chimeric peptide, GEP44, on core temperature in male and female DIO GLP-1R^+/+^ and GLP-1R^-/-^ mice. Mice were maintained on HFD (60% kcal from fat; N=8-9/group) for approximately 4 months prior to receiving SC injections of vehicle (sterile saline/water) followed by escalating doses of GEP44 (5, 10, 20 and 50 nmol/kg; 3 mL/kg injection volume). *A*, Effect of GEP44 on change in core temperature in male HFD-fed DIO GLP-1R^+/+^ mice; *B*, Effect of GEP44 on change in core temperature in male HFD-fed DIO GLP-1R ^-/-^ mice; *C*, Effect of GEP44 on change in core temperature in female HFD-fed DIO GLP-1R^+/+^ mice; *D*, Effect of GEP44 on change in core temperature in female HFD-fed DIO GLP-1R^-/-^ mice. Data are expressed as mean ± SEM. \**P*<0.05 GEP44 vs. vehicle.

**Figure 6A-D:**
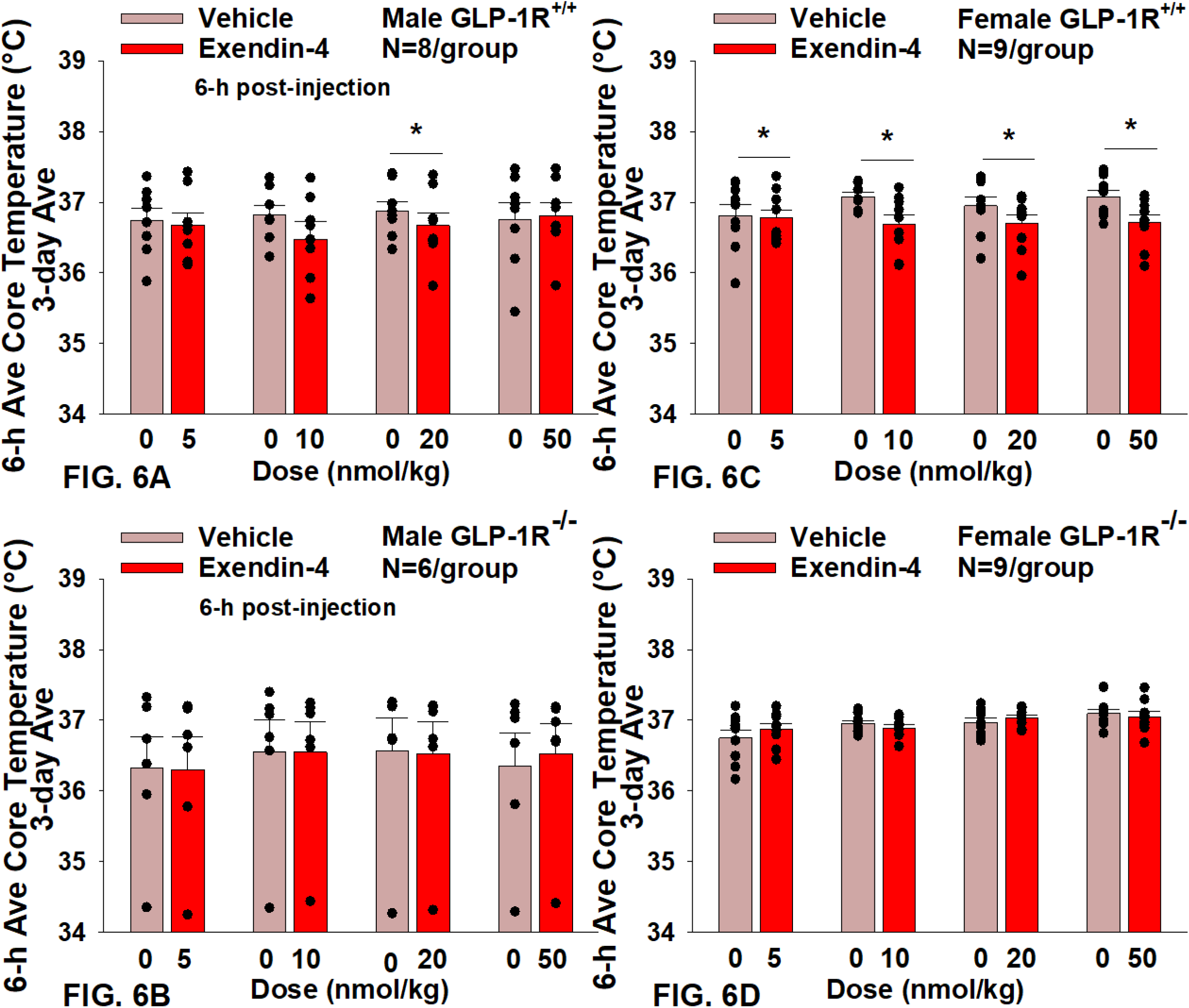
Effects of the selective GLP-1R agonist, exendin-4, on core temperature in male and female DIO GLP-1R^+/+^ and GLP-1R^-/-^ mice. Mice were maintained on HFD (60% kcal from fat; N=6-9/group) for approximately 4 months prior to receiving SC injections of vehicle (sterile saline/water) followed by escalating doses of GEP44 (5, 10, 20 and 50 nmol/kg; 3 mL/kg injection volume). *A*, Effect of exendin-4 on change in core temperature in male HFD-fed DIO GLP-1R^+/+^ mice; *B*, Effect of exendin-4 on change in core temperature in male HFD-fed DIO GLP-1R^-/-^ mice; *C*, Effect of exendin-4 on change in core temperature in female HFD-fed DIO GLP-1R^+/+^ mice; *D*, Effect of exendin-4 on change in core temperature in female HFD-fed DIO GLP-1R^-/-^ mice. Data are expressed as mean ± SEM. \**P*<0.05 exendin-4 vs. vehicle.

#### Activity

GEP44 and exendin-4 both reduced activity over 6 hours post-injection in male (**Fig. 7A**; *P*<0.05) and female DIO GLP-1R^+/+^ mice (**Fig. 7C**; *P*<0.05). Similar results also observed over the 12-h dark cycle in response to GEP44 in male and female DIO GLP-1R^+/+^ mice (*P*<0.05; data not shown). However, the effect of GEP44 to reduce activity was observed only at the low and high dose in male GLP-1R^-/-^ mice (**Fig. 7B**; *P*<0.05) and not in female GLP-1R^-/-^ mice (**Fig. 7D**; *P*=NS). There was an effect of exendin-4 to reduce activity in male (**Fig. 8A**; *P*<0.05) and female GLP-1R^+/+^ mice (**Fig. 8C**; *P*=NS) but not in GLP-1R^-/-^ mice (**Fig. 8B**; **Fig. 8D**; *P*=NS).

**Figure 7A-D.**
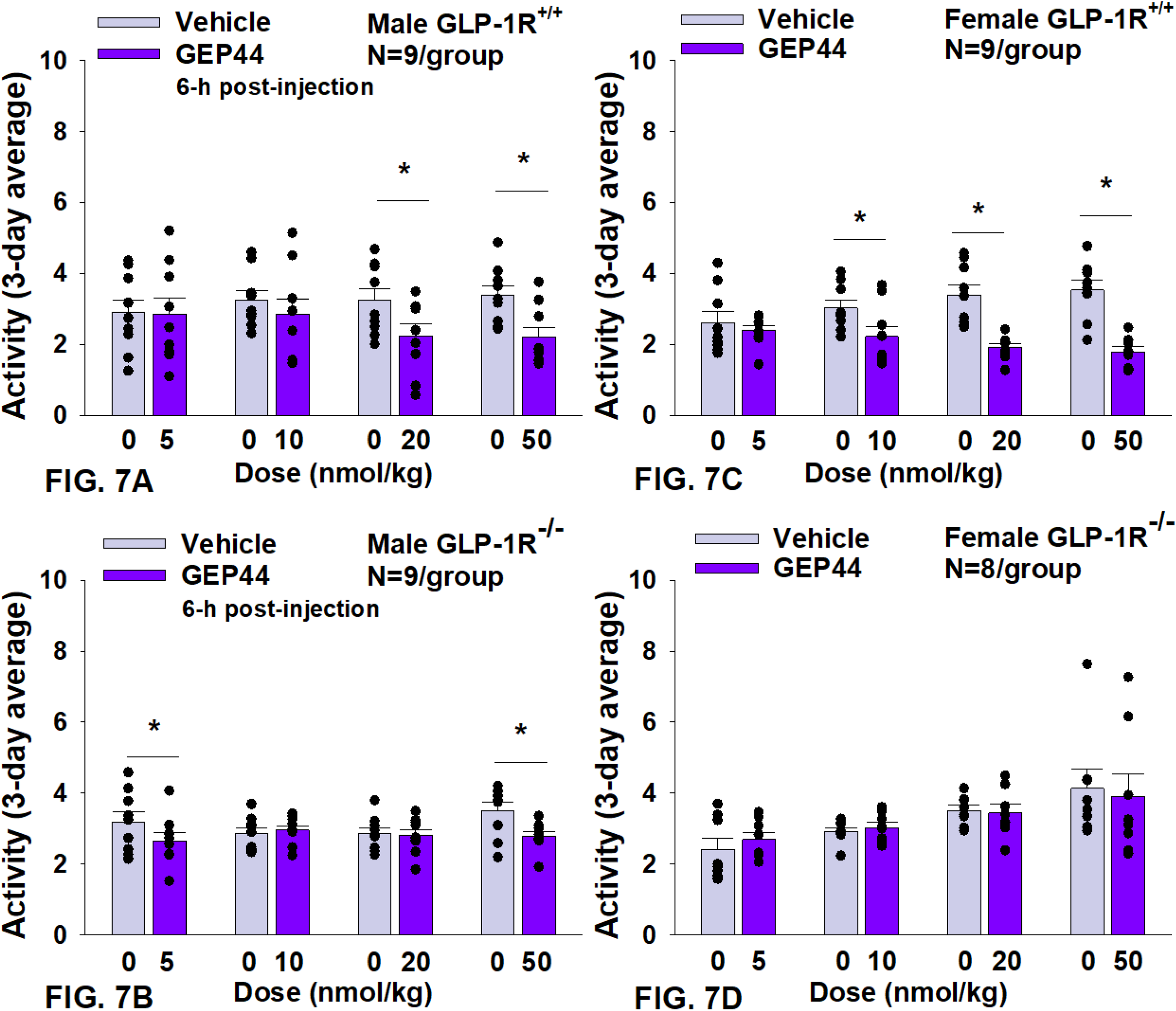
Effects of the chimeric peptide, GEP44, on activity in male and female DIO GLP-1R^+/+^ and GLP-1R^-/-^ mice. Mice were maintained on HFD (60% kcal from fat; N=7-9/group) for approximately 4 months prior to receiving SC injections of vehicle (sterile saline/water) followed by escalating doses of GEP44 (5, 10, 20 and 50 nmol/kg; 3 mL/kg injection volume). *A*, Effect of GEP44 on change in activity in male HFD-fed DIO GLP-1R^+/+^ mice; *B*, Effect of GEP44 on change in activity in male HFD-fed DIO GLP-1R^-/-^ mice; *C*, Effect of GEP44 on change in activity in female HFD-fed DIO GLP-1R^+/+^ mice; *D*, Effect of GEP44 on change in activity in female HFD-fed DIO GLP- 1R^-/-^ mice. Data are expressed as mean ± SEM. \**P*<0.05 GEP44 vs. vehicle.

**Figure 8A-D:**
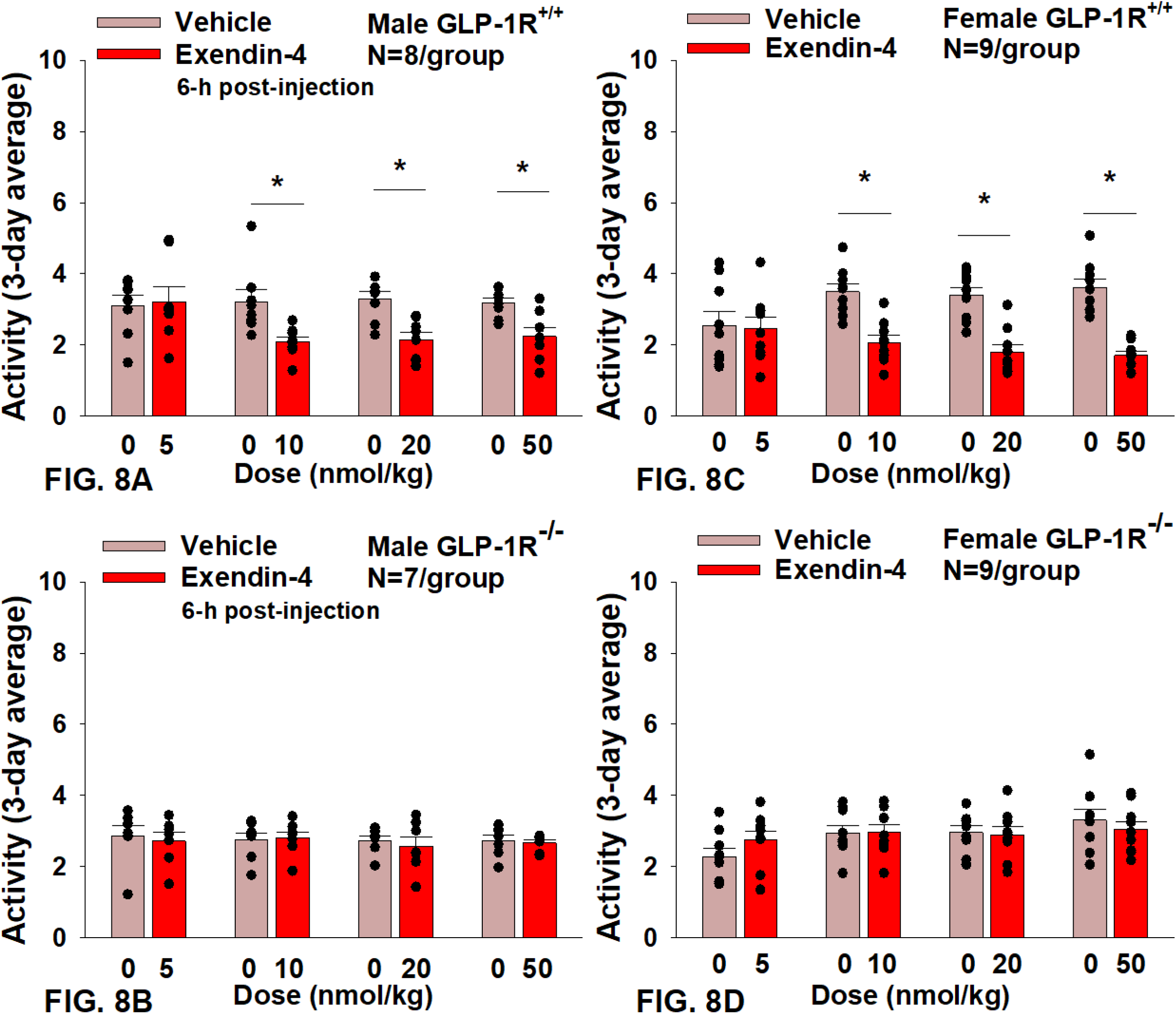
Effects of the selective GLP-1R agonist, exendin-4, on activity in male and female DIO GLP-1R^+/+^ and GLP-1R^-/-^ mice. Mice were maintained on HFD (60% kcal from fat; N=8-9/group) for approximately 4 months prior to receiving SC injections of vehicle (sterile saline/water) followed by escalating doses of GEP44 (5, 10, 20 and 50 nmol/kg; 3 mL/kg injection volume). *A*, Effect of exendin-4 on change in activity in male HFD-fed DIO GLP-1R^+/+^ mice; *B*, Effect of exendin-4 on change in activity in male HFD-fed DIO GLP-1R^-/-^ mice; *C*, Effect of exendin-4 on change in activity in female HFD-fed DIO GLP-1R^+/+^ mice; *D*, Effect of exendin-4 on change in activity in female HFD-fed DIO GLP-1R^-/-^ mice. Data are expressed as mean ± SEM. \**P*<0.05 exendin-4 vs. vehicle.

### Tissue Collection for Quantitative Real-Time PCR (qPCR)

As an additional readout of GEP44 and exendin-4-elicited thermogenic effects in IBAT, relative levels of mRNA for UCP-1, Gpr120, Ppargc1a, Cidea, and Dio2 were compared by PCR in response to GEP44 (50 nmol/kg) and exendin-4 (50 nmol/kg) or vehicle treatment at 2-h post-injection in male (**Table 1A**) and female GLP-1R^+/+^ mice (**Table 1C**). We found that GEP44 increased Ppargc1a in male GLP-1R^+/+^ mice [F(1,20) = 7.506, *P*=0.013]. Similarly, GEP44 increased Ppargc1a in female GLP-1R^+/+^ mice [F(1,18) = 36.773, P<0.001]. In contrast, the effects of GEP44 on thermogenic gene expression were largely absent in male GLP-1R^-/-^ mice. In addition, GEP44 stimulated GPR120 in female GLP-1R^-/-^ mice.

In female GLP-1R^+/+^ mice, exendin-4 increased UCP-1 [F(1,18) = 5.967, *P*=0.025], Gpr120 [F(1,18) = 10.744, *P*=0.004], and Ppargc1a [F(1,18) = 19.767, *P*<0.001] while the effects of exendin-4 on thermogenic gene expression were absent in male and female GLP-1R^-/-^ mice.

We also implanted a subset of mice with temperature transponders (HTEC IPTT-300; BIO MEDIC DATA SYSTEMS, INC, Seaford, DE) underneath both IBAT pads in order to obtain a more functional measure of IBAT thermogenesis (IBAT temperature (TIBAT)) as previously described [16; 18]. Similar to what we found with GEP44-elicited increases in IBAT thermogenic gene expression, we found that GEP44 (50 nmol/kg) also increased TIBAT at 240-min post-injection in male DIO GLP-1R^+/+^ mice (N=3/group; *P*<0.05) (**Figure 9A**). Likewise, exendin-4 (50 nmol/kg) also increased TIBAT at 180 and 360-min post-injection (N=4/group; *P*<0.05) (**Figure 9B**).

**Figure 9A-B:**
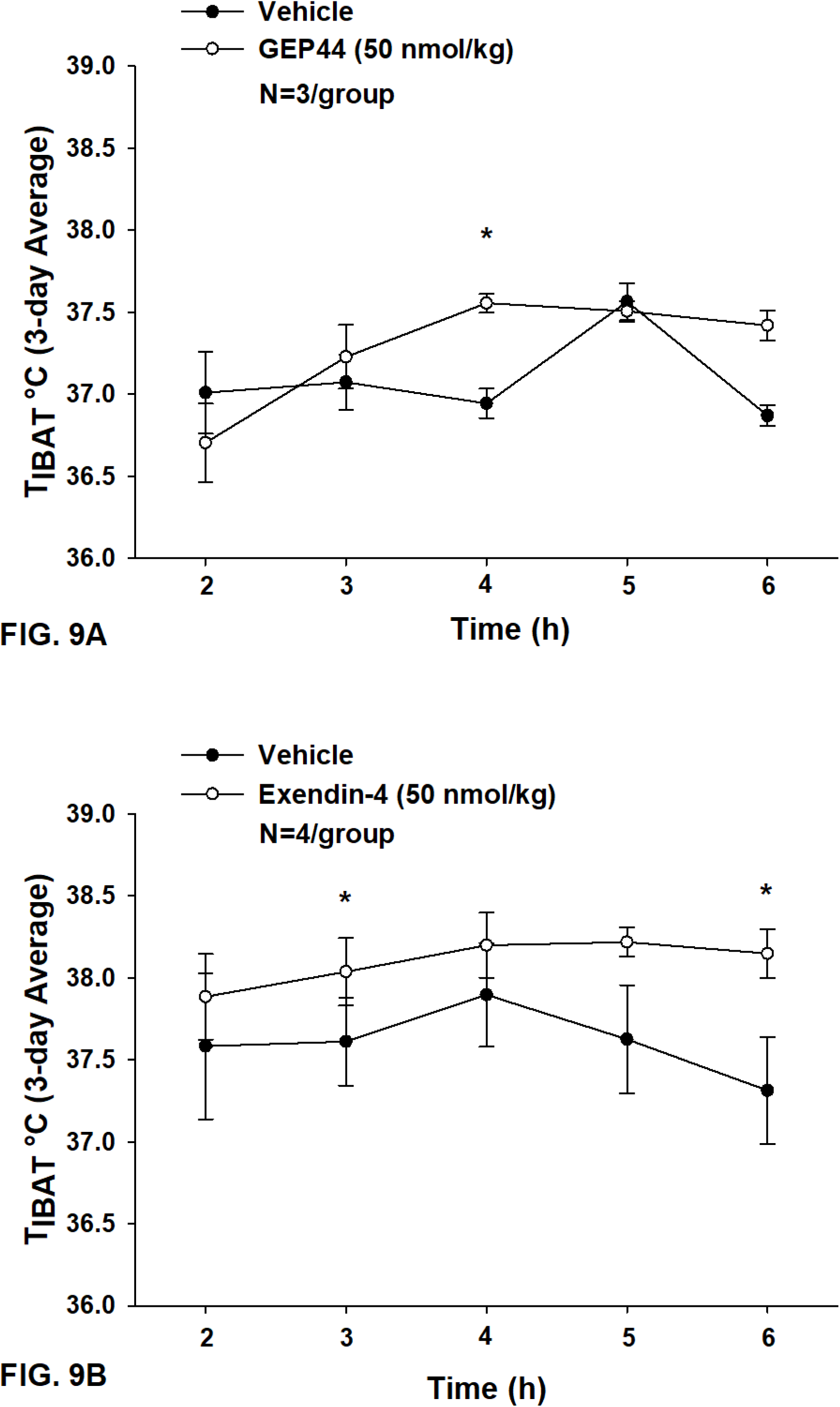
Effects of GEP44 and exendin-4 on IBAT temperature (TIBAT) in male DIO GLP-1R^+/+^ mice. *A*, Effect of GEP44 on TIBAT in male DIO GLP-1R^+/+^ mice; *B*, Effect of exendin-4 on TIBAT in male DIO GLP-1R^+/+^ mice. Data are expressed as mean ± SEM. \**P*<0.05 GEP44 or exendin-4 vs. vehicle.

Lower doses of exendin-4 largely reproduced the effects found at the higher dose (50 nmol/kg). Namely, exendin-4 (10 nmol/kg) tended to increase TIBAT at 300-min post-injection (*P*<0.05; data not shown) while the slightly higher dose (20 nmol/kg) increased TIBAT at 240-min post-injection (*P*<0.05; data not shown). Exendin-4 (20 nmol/kg) appeared to produce a reduction of TIBAT at 120-min post-injection (*P*<0.05; data not shown), but this was not observed at other doses. In contrast, GEP44 failed to produce significant effects on TIBAT at lower doses (data not shown).

Together, our findings suggest that while both GEP44 and exendin-4 increase BAT thermogenesis, the effects of exendin-4 on BAT thermogenesis (TIBAT) appeared to be longer lasting relative to GEP44.

### Blood Glucose and Plasma Hormones

Consistent with previous findings in rats, we found that that GEP44 also reduced tail vein glucose in both male (**Figure 10A**) [F(1,14) = 39.938, *P*<0.05)] and female GLP- 1R^+/+^ DIO mice (**Figure 10B**) [F(1,10) = 6.954, *P*<0.05]. Moreover, the effects of GEP44 to reduce tail vein glucose were absent in both male [F(1,5) = 28.122, *P*<0.05] and female GLP-1R^-/-^ mice [F(1,2) = 28.189, *P*<0.05]. In addition, exendin-4 reduced fasting tail vein glucose in both male (**Figure 10B**) [F(1,14) = 41.690, *P*<0.05] and female GLP-1R^+/+^ DIO mice (**Figure 10B**) [F(1,9) = 7.241, *P*<0.05]. As was the case with GEP44, the effects of exendin-4 to reduce tail vein blood glucose were also blocked in male GLP-1R^-/-^ mice [F(1,4) = 54.475, *P*<0.05]. The effect of exendin-4 to reduce glucose also appeared to be impaired in female GLP-1R^-/-^ mice [F(1,1) = 78.797, *P*=0.071]. In addition, exendin-4 reduced plasma leptin in male DIO mice [F(1,12) = 8.522, *P*<0.05] (**Table 3A**) and tended to reduce leptin in female DIO mice [F(1,15) = 4.157, *P*=0.059] (**Table 3B**). In addition, GEP44 tended to reduce plasma leptin in female DIO mice [F(1,14) = 3.325, *P*=0.09] **(Table 3B)**. Both GEP44 [F(1,14) = 5.405, P<0.05] and exendin-4 [F(1,15) = 5.151, *P*<0.05] also reduced total cholesterol in female DIO mice (**Table 3B**). These effects on total cholesterol also tended to be observed in response to both exendin-4 [F(1,14) = 5.169, *P*<0.05] and GEP44 [F(1,13) = 3.805, *P*=0.073] in female GLP-1R^-/-^ mice (**Table 3B**).

**Figure 10A-B:**
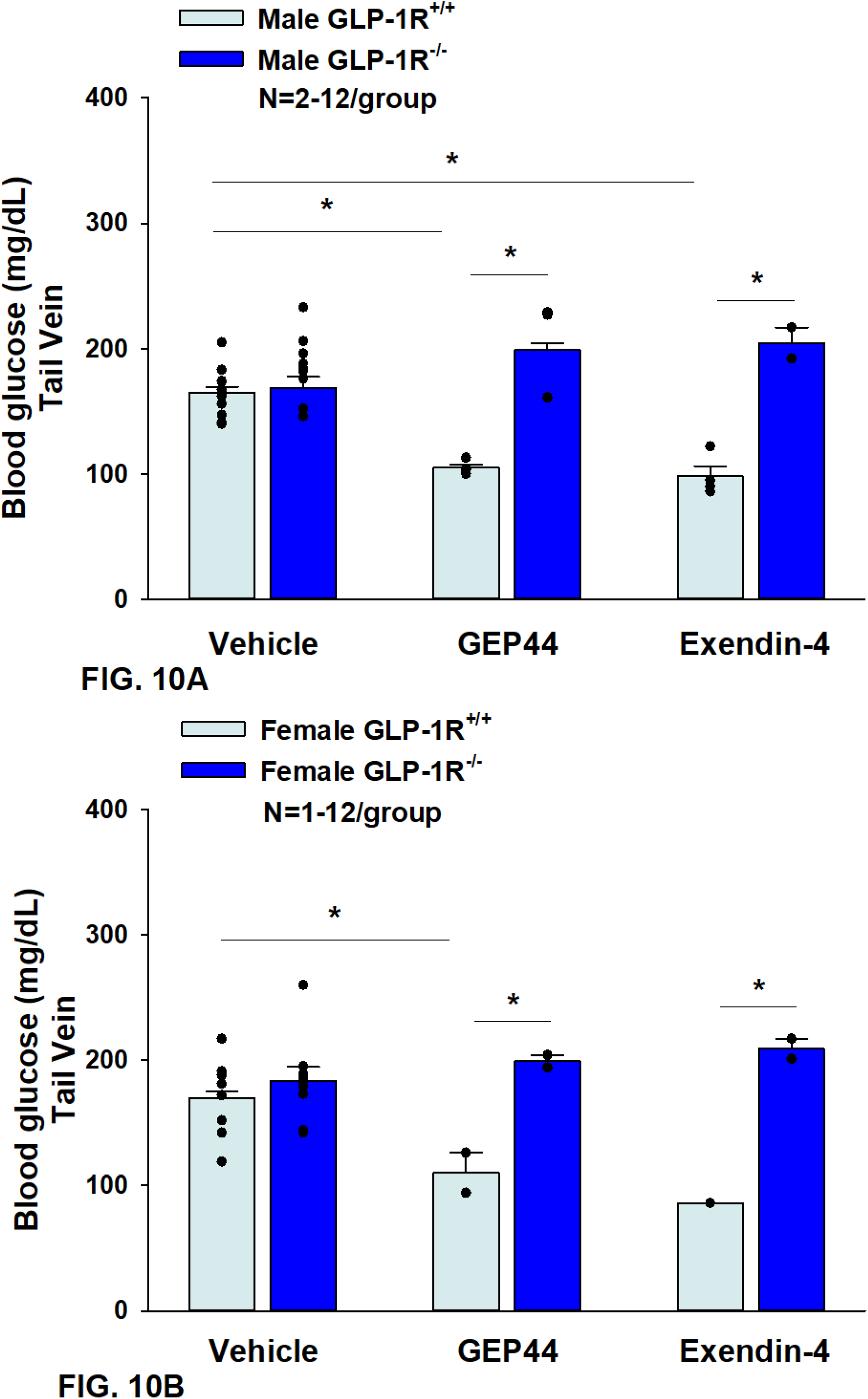
Effects of GEP44 and exendin-4 on tail vein glucose in male and female DIO GLP-1R^+/+^ and GLP-1R^-/-^ mice. *A*, Effect of GEP44 and exendin-4 on tail vein glucose in male DIO GLP-1R^+/+^ and GLP-1R^-/-^ mice; *B*, Effect of GEP44 and exendin-4 on tail vein glucose in female DIO GLP-1R^+/+^ and GLP-1R^-/-^ mice. Blood was collected by tail vein nick (glucose) at 2-h post-injection of VEH, exendin-4 (50 nmol/kg) or GEP44 (50 nmol/kg). Data are expressed as mean ± SEM. \**P*<0.05 GEP44 or exendin-4 vs. vehicle.

**Table 2A-D.**
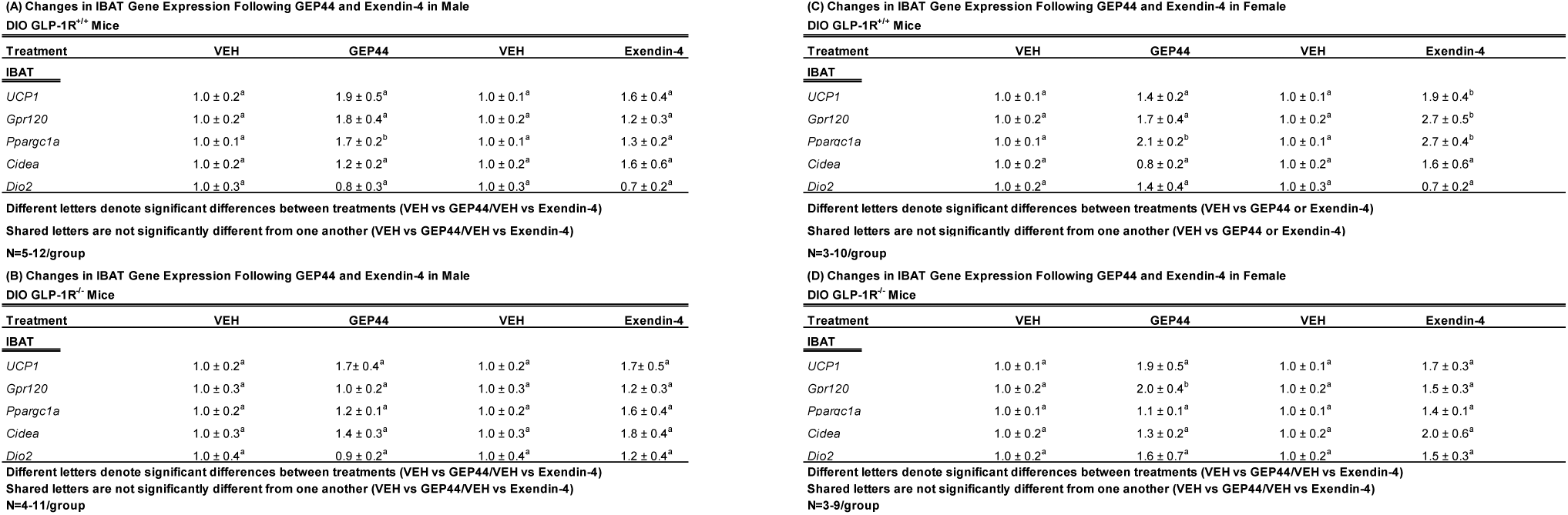
Gene expression in IBAT following SC vehicle, GEP44 (50 nmol/kg) or exendin-4 (50 nmol/kg) treatment in male and female GLP-1R^+/+^ and GLP-1R^-/-^ DIO mice (N=7-12/group). Data are expressed as mean ± SEM. \**P*<0.05 GEP44 or exendin- 4 vs. vehicle.

**Table 3A-B:**
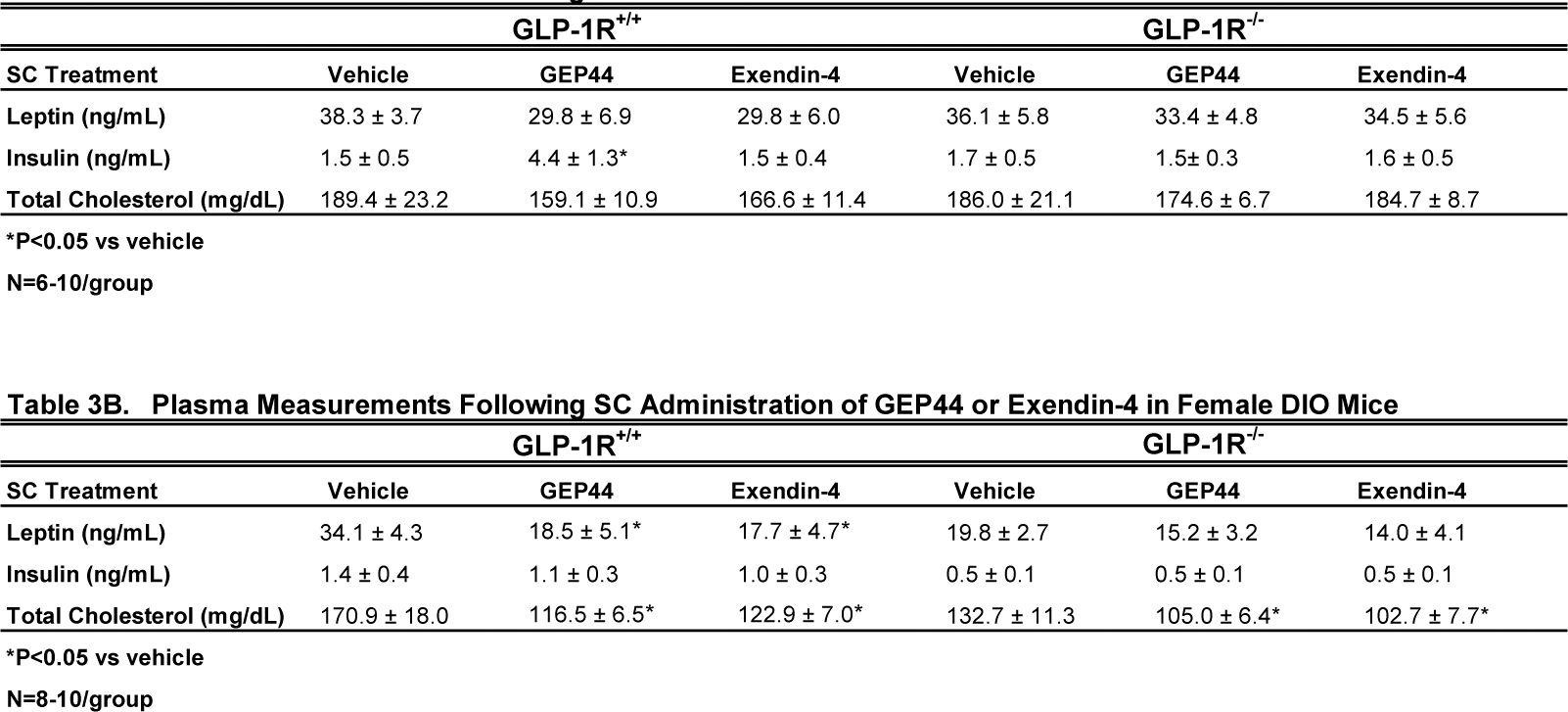
Effects of the chimeric peptide, GEP44 (50 nmol/kg) and the selective GLP-1R agonist, exendin-4, on plasma leptin, insulin, and total cholesterol in A) male and B) female DIO GLP-1R^+/+^ and GLP-1R^-/-^ mice (N=7-12/group). Blood was collected by tail vein nick (glucose) or cardiac stick (leptin, insulin, total cholesterol) at 2-h post-injection of VEH, exendin-4 (50 nmol/kg) or GEP44 (50 nmol/kg). Data are expressed as mean ± SEM. \**P*<0.05 GEP44 or exendin-4 vs. vehicle.

## Discussion

Here we report effects of the novel chimeric peptide (GEP44), which targets GLP-1R, Y1R and Y2R, on energy intake, BW, thermoregulation (core temperature) and gross motor activity in DIO mice and characterize the extent to which these effects are mediated through GLP-1R. We tested the hypothesis that GEP44 reduces energy intake and BW through a GLP-1R dependent mechanism. We found that GEP44 reduced BW in both male and female DIO male GLP-1R^+/+^ mice whereas these effects were absent in male and female DIO GLP-1R^-/-^ mice. These findings suggest that GLP- 1R signaling contributes to GEP44-elicited reduction of BW in both male and female mice. Additionally, GEP44 decreased energy intake in both male and female DIO GLP- 1R^+/+^ mice, but GEP44 appeared to produce more consistent effects across multiple doses in males. These findings suggest that 1) GEP44 reduces body weight, in part, through reductions in energy intake in DIO mice and 2) male mice might have enhanced sensitivity to the anorexigenic effects of GEP44. In GLP-1R^-/-^ mice, the effects on energy intake were observed only at low and high doses in males suggesting that GEP44 may reduce energy intake in males, in part, through a GLP-1R independent mechanism. GEP44 reduced both core temperature and activity in both male and female GLP-1R^+/+^ mice suggesting that reductions in energy expenditure and/or spontaneous activity-induced thermogenesis may also contribute to the weight lowering effects of GEP44 in mice. Lastly, we show that GEP44 reduced fasting blood glucose in DIO male and female mice through GLP-1R signaling. Together, these findings support the hypothesis that the chimeric peptide, GEP44, reduces energy intake, BW, core temperature, and glucose levels, in part, through a GLP-1R dependent mechanism.

We extend previous findings from our laboratory [12; 13] and demonstrate that the effects of GEP44 to elicit weight loss in both male and female mice are primarily driven by GLP-1R. These effects are mediated, at least in part, by reductions of energy intake. Similar to GLP-1R driven effects of GEP44 on BW, based on our findings in both male and female GLP-1R^-/-^ mice, the ability of GEP44 to reduce energy intake also appears to be largely mediated by GLP-1R. However, the finding that the low and high dose of GEP44 also reduced energy intake in male GLP-1R^-/-^ mice suggests other mechanisms are involved in contributing to these effects. Given the role of Y2R in the control of energy intake and BW [20], and results showing synergism/additive effects between co-administered GLP-1R and Y2-R agonists [21; 22], it is likely that the Y2R is also playing a role in the beneficial effects of GEP44 in terms of BW.

We incorporated the use of exendin-4 as a positive control for GLP-1R mediated effects on both food intake and body weight. Similar to what others have reported [14; 23], we also report that exendin-4 reduced BW and energy intake in both male and female DIO mice through GLP-1R signaling. These findings are consistent with previous findings from Baggio and colleagues who reported that a single dose of exendin-4 (1.5 μg or 0.356 nmol/mouse) failed to reduce chow diet intake in male GLP-1R^-/-^ mice [14]. Another study also found that chronic subcutaneous administration of a single dose (0.126 mg/kg/day or 30 nmol/kg/day) also failed to reduce high fat diet intake and BW (vehicle corrected) in male GLP-1R^-/-^ mice [23]. In contrast to our studies, only a single dose of exendin-4 was examined in male mice in both studies [14; 23] and neither study examined the effects of exendin-4 in male and female DIO mice.

We also extend previous findings from our laboratory [12; 13] to demonstrate that GEP44 reduces core temperature (surrogate measure of energy expenditure) in both male and female mice, similar to what we described earlier herein following exendin-4 treatment. These data are also consistent with a previous report by Hayes and colleagues showing that the GLP-1R agonist, exendin-4, produced a long-lasting reduction of core temperature (hypothermia) that lasted for 4 hours following systemic (intraperitoneal (IP)) injections in rats [24]. The hypothermic effects may be due, in part, to the reduced thermic effect of food (diet-induced thermogenesis) in exendin-4 and GEP44 treated mice. Furthermore, Baggio and colleagues [14] found that central and peripheral (IP; 1.5 μg or 0.356 nmol/mouse) administration of exendin-4 reduced VO2 and resting energy expenditure in adult male wild-type mice. These effects occurred over 2 and 21 hours following peripheral and central administration, respectively.

Similarly, Krieger and colleagues found that peripheral exendin-4 reduced energy expenditure over 4-h post-injection in rats [25]. In addition, van Eyk and colleagues found that liraglutide treatment reduced resting energy expenditure after 4 weeks of treatment in humans with type 2 diabetes [26]. Gabery initially found that semaglutide reduced energy expenditure during the dark phase in DIO mice through treatment day 6 but these differences were no longer significant after adjusting for lean mass [27].

Similarly, Blundell and colleagues initially found that semaglutide reduced resting energy expenditure after 12 weeks of treatment in humans but these effects were no longer significant after adjusting for lean mass [28]. Despite previous reports indicating that acute central (ICV) administration of GLP-1 increases core temperature over 4-h post-injection [29], IBAT temperature over 6 hours post-injection [30] and activates SNS fibers that innervate BAT [30], to our knowledge, the majority of studies that administered GLP-1R agonists peripherally have either found a reduction or no change in energy expenditure. Given the dearth of studies that have measured both core temperature and energy expenditure in the same group of animals, it will be important to measure both endpoints simultaneously in the same animal model. Ongoing studies are currently in the process of addressing the extent to which systemic administration of GEP44 and exendin-4 alters core temperature and more directly impacts energy expenditure (measured by indirect calorimetry) in male and female DIO rats.

In contrast to the effects of GEP44 and exendin-4 to reduce core temperature, we found that both GEP44 and exendin-4 stimulated thermogenic markers (biochemical readout of BAT thermogenesis) within IBAT and TIBAT (functional readout of BAT thermogenesis). Our findings, however, that exendin-4 stimulated thermogenic markers within IBAT and IBAT temperature is consistent with what others have found following systemic (intraperitoneal) exendin-4 [31] or liraglutide [32] administration in mice. On the other hand, Krieger [25] found that systemic (IP) exendin-4 reduced BAT thermogenesis, skin temperature above the interscapular area over 2-h post-injection and BAT Adrb3 in a rat model. Note that measurements of skin temperature above the interscapular area might not necessarily reflect TIBAT as the interscapular subcutaneous area may also be influenced by heating and cooling of the skin temperature. We found that the majority of thermogenic genes in IBAT were elevated in response to both GEP44 and exendin-4. Furthermore, our findings suggest that GEP44 may also elicit BAT thermogenesis through similar mechanisms (PPargc1a) in both male and female GLP-1R^+/+^ mice while exendin-4 may increase BAT thermogenesis via multiple thermogenic genes (UCP-1, Gpr120, and Ppargc1a) in female mice. In contrast to the effects of exendin-4 in GLP-1R^+/+^ mice, the chimeric peptide, GEP44, appears to promote BAT thermogenesis in both male and female GLP-1R^+/+^ mice through different genes. In contrast, GEP44 stimulated GPR120 only in female GLP-1R^-/-^ mice raising the possibility that other receptor subtypes may contribute to these effects at the high dose in female mice. Overall, our findings suggest that systemic administration of both GEP44 and exendin-4 may promote BAT thermogenesis through different mechanisms in a mouse model. It will be important in future studies to examine if SNS outflow to BAT is a predominant mediator of GEP44 or exendin-4-elicited weight loss in male and female DIO mice.

Our findings also demonstrate that GEP44 reduced gross motor activity through what appears to be both GLP-1R and Y2R signaling in male mice; while this effect appears to be entirely mediated by GLP-1R in female mice. In contrast, we found that exendin-4 treatment reduces activity strictly through GLP-1R in both male and female mice, as anticipated. Similarly, others have also found that exendin-4 at lower [0.5-5 μg/kg (0.2-2 μg/rat, IP)] [33] or higher doses [30 μg/kg (9 μg/rat, IP)] [34] reduces locomotor activity in lean male Sprague Dawley rats [33; 34]. While Hayes and colleagues [24] did not find any change in activity in response to exendin-4 at [3 μg/kg (0.9 μg/rat, IP)] in lean male Sprague-Dawley rats, it might be possible that differences in paradigm and/or timing of injections may account for discrepancies between studies. Given that we found reductions in both core temperature and gross motor activity, our data raise the possibility that reductions in spontaneous physical activity-induced thermogenesis [35] and/or shivering and non-shivering thermogenesis in skeletal muscle [36] may have contributed to the hypothermic effects of both GEP44 and exendin-4. Moreover, our findings indicate that GEP44 may produce GLP-1R independent effects on 1) activity in male mice and 2) core temperature in both male and female mice while exendin-4 appears to impact both activity and core temperature through GLP-1R dependent mechanism in both male and female mice.

We have previously found that GEP44 improves glucoregulation in a rat model [13]. Specifically, GEP44 improves fasting blood glucose and glucose tolerance in DIO rats [13]. These findings are consistent with the weight lowering and/or glucose lowering effects of peripheral GEP44 [12; 13] and exendin-4 [31] in rats and mice, respectively. In addition, GEP44 was found to improve glucose tolerance in a mammalian model (musk shrew) [13]. We have now extended these findings by showing that the effects of GEP44 to reduce fasting blood glucose in DIO male and female mice were mediated, in part, by GLP-1R. We acknowledge that GEP44- and exendin-4 elicited effects on fasting blood glucose may also be due, in part, to GEP44-elicited weight loss and future studies will include weight-restricted controls. In addition, the effects of peripheral GEP44 and exendin-4 to reduce cholesterol in female GLP-1R^+/+^ and GLP-1R^-/-^ mice are also consistent with the potential GLP-1R and/or Y2-receptor mediated cholesterol lowering effects observed following the GLP-1R agonists, liraglutide [37] and exendin-4 [38; 39], Y2 receptor agonist, lapidated PYY3-36 analog, [Lys7(C16-γGlu)PYY3−36] (similar potency on Y2R as PYY3-36) [37], and dual GLP-1R/Y2 receptor agonists, 6q and peptide 19 [37], in DIO rodents.

One limitation to our studies is the lack of weight restricted or pair-fed controls for studies that assessed the effects of GEP44 and exendin-4 on gene expression as well as plasma measurements (total cholesterol, leptin, and glucose) in adult DIO male and female mice. As a result, it is possible that the effects of GEP44 and exendin-4 on thermogenic gene expression or glucose, total cholesterol or leptin lowering in DIO mice may also be due, in part, to the weight loss in response to GEP44 or exendin-4 treatment. In addition, we only collected TIBAT between 120 and 360 min-post-injection. It is possible that, due to absence of being able to measure TIBAT at earlier time points, we might have missed any potential hypothermic effects that might have preceded the elevations of TIBAT that occurred at the later time points. In addition, we were unable to assess the roles of Y1R or Y2R signaling more fully in contributing to the effects of GEP44. Future studies incorporating the use of Y2R deficient mice will be helpful in more fully establishing the role of Y2R signaling in the anorectic response to GEP44.

In summary, the results presented in this manuscript highlight the beneficial effects of the dual-agonist, GEP44, on BW, energy intake, and glucoregulation in adult male and female DIO mice at doses that have not been found to elicit visceral illness in rats or emesis in shrews [12; 13]. Our findings further demonstrate that effects on changes of BW, energy intake, activity, and glucose levels are largely mediated through GLP-1R signaling. However, given that GEP44 may also elicit GLP-1R independent effects on 1) activity in male mice and 2) core temperature in both male and female mice, it is possible that other receptor subtypes, including the Y1R [40] and Y2R, may contribute to these effects. Given the positive findings of GEP44 in male and female DIO mice and rats [12; 13], it will be important to examine the safety and effectiveness of long-term treatment of GEP44 on BW and cardiovascular function in obese nonhuman primates prior to trials in humans. The findings to date indicate that GEP44 is a promising drug targeting limitations associated with current GLP-1R agonist medications [41; 42] for the treatment of obesity and/or T2DM.

## ACKNOWLEDGMENTS

Plasma sample analysis was performed by the UC Davis MMPC Live M&MH Core (NIH U2CDK135074). The authors thank the technical support of Dr. Tami Wolden-Hanson with body composition measurements and James Graham and Dr. Peter Havel for plasma analyses.

## Figure legend

**Supplemental Figure 1A-D:**
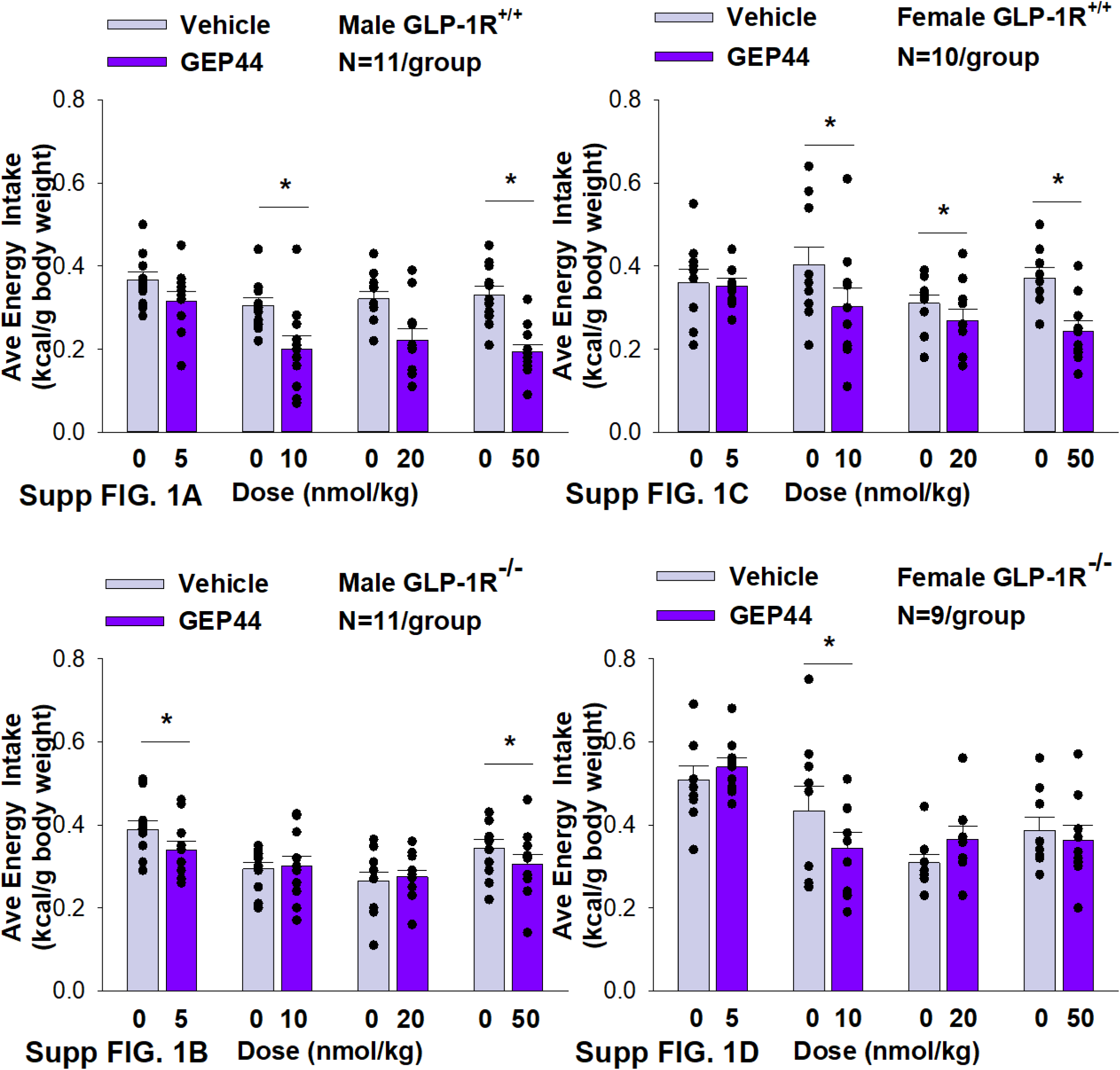
Effects of the chimeric peptide, GEP44, on energy intake (kcal/gram BW) in male and female DIO GLP-1R^+/+^ and GLP-1R^-/-^ mice. Mice were maintained on HFD (60% kcal from fat; N=10-11/group) for approximately 4 months prior to receiving SC injections of vehicle (sterile saline/water) followed by escalating doses of GEP44 (5, 10, 20 and 50 nmol/kg; 3 mL/kg injection volume). *A*, Effect of GEP44 on energy intake in male HFD-fed DIO GLP-1R^+/+^ mice; *B*, Effect of GEP44 on change on energy intake in male HFD-fed DIO GLP-1R^-/-^ mice; *C*, Effect of GEP44 on energy intake in male HFD-fed DIO GLP-1R^+/+^ mice; *D*, Effect of GEP44 on energy intake in male HFD-fed DIO GLP-1R ^-/-^ mice. Data are expressed as mean ± SEM. \**P*<0.05 GEP44 vs. vehicle.

**Supplemental Figure 2A-D:**
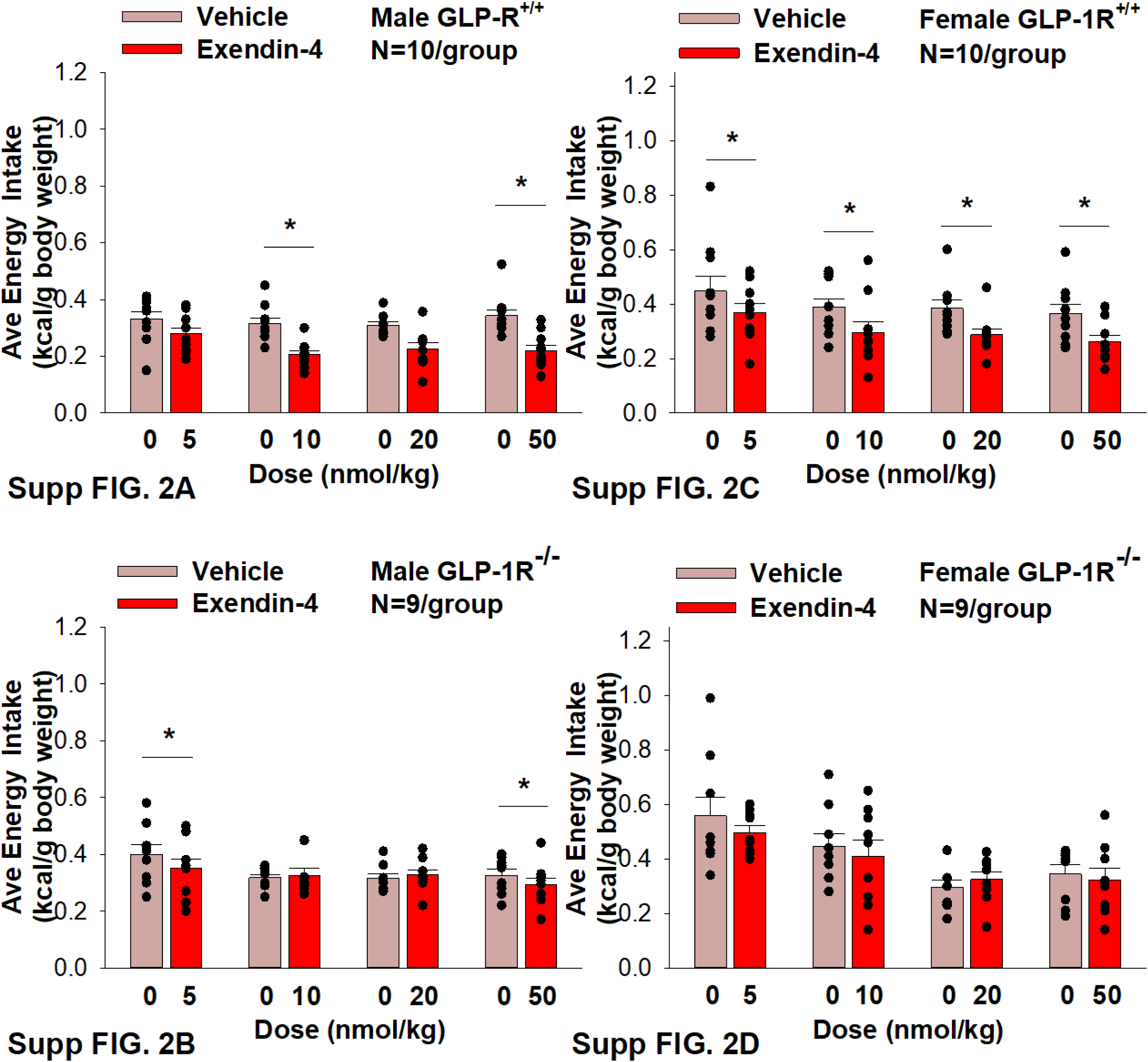
Effects of the selective GLP-1R agonist, exendin-4, on energy intake (kcal/gram BW) in male and female DIO GLP-1R^+/+^ and GLP-1R^-/-^ mice. Mice were maintained on HFD (60% kcal from fat; N=9-11/group) for approximately 4 months prior to receiving SC injections of vehicle followed by escalating doses of GEP44 (5, 10, 20 and 50 nmol/kg; 3 mL/kg injection volume). *A*, Effect of GEP44 on energy intake in male HFD-fed DIO GLP-1R^+/+^ mice; *B*, Effect of GEP44 on change on energy intake in male HFD-fed DIO GLP-1R^-/-^ mice; *C*, Effect of GEP44 on energy intake in male HFD-fed DIO GLP-1R^+/+^ mice; *D*, Effect of GEP44 on energy intake in male HFD-fed DIO GLP-1R^-/-^ mice. Data are expressed as mean ± SEM. \**P*<0.05 exendin-4 vs. vehicle.

## References

[1] N.C.D.R.F. Collaboration, Worldwide trends in underweight and obesity from 1990 to 2022: a pooled analysis of 3663 population-representative studies with 222 million children, adolescents, and adults. Lancet 403 (2024) 1027–1050.

[2] Z.J. Ward, S.N. Bleich, A.L. Cradock, J.L. Barrett, C.M. Giles, C. Flax, M.W. Long, and S.L. Gortmaker, Projected U.S. State-Level Prevalence of Adult Obesity and Severe Obesity. The New England journal of medicine 381 (2019) 2440–2450.

[3] A. Okunogbe, R. Nugent, G. Spencer, J. Powis, J. Ralston, and J. Wilding, Economic impacts of overweight and obesity: current and future estimates for 161 countries. BMJ Glob Health 7 (2022).

[4] J.P.H. Wilding, R.L. Batterham, S. Calanna, M. Davies, L.F. Van Gaal, I. Lingvay, B.M. McGowan, J. Rosenstock, M.T.D. Tran, T.A. Wadden, S. Wharton, K. Yokote, N. Zeuthen, R.F. Kushner, and S.S. Group, Once-Weekly Semaglutide in Adults with Overweight or Obesity. The New England journal of medicine 384 (2021) 989.

[5] R.J. Rodgers, M.H. Tschop, and J.P. Wilding, Anti-obesity drugs: past, present and future. Disease models & mechanisms 5 (2012) 621–6.

[6] H.L. Daneschvar, M.D. Aronson, and G.W. Smetana, FDA-Approved Anti-Obesity Drugs in the United States. Am J Med 129 (2016) 879 e1–6.

[7] D.B. Allison, K.M. Gadde, W.T. Garvey, C.A. Peterson, M.L. Schwiers, T. Najarian, P.Y. Tam, B. Troupin, and W.W. Day, Controlled-release phentermine/topiramate in severely obese adults: a randomized controlled trial (EQUIP). Obesity 20 (2012) 330–42.

[8] C.M. Apovian, L. Aronne, D. Rubino, C. Still, H. Wyatt, C. Burns, D. Kim, E. Dunayevich, and C.-I.S. Group, A randomized, phase 3 trial of naltrexone SR/bupropion SR on weight and obesity-related risk factors (COR-II). Obesity 21 (2013) 935–43.

[9] A.M. Jastreboff, L.J. Aronne, N.N. Ahmad, S. Wharton, L. Connery, B. Alves, A. Kiyosue, S. Zhang, B. Liu, M.C. Bunck, A. Stefanski, and S.-. Investigators, Tirzepatide Once Weekly for the Treatment of Obesity. The New England journal of medicine 387 (2022) 205–216.

[10] L.J. Aronne, N. Sattar, D.B. Horn, H.E. Bays, S. Wharton, W.Y. Lin, N.N. Ahmad, S. Zhang, R. Liao, M.C. Bunck, I. Jouravskaya, M.A. Murphy, and S.-. Investigators, Continued Treatment With Tirzepatide for Maintenance of Weight Reduction in Adults With Obesity: The SURMOUNT-4 Randomized Clinical Trial. JAMA : the journal of the American Medical Association (2023).

[11] A.M. Jastreboff, L.M. Kaplan, and M.L. Hartman, Triple-Hormone-Receptor Agonist Retatrutide for Obesity. Reply. The New England journal of medicine 389 (2023) 1629–1630.

[12] K.S. Chichura, C.T. Elfers, T.S. Salameh, V. Kamat, O.G. Chepurny, A. McGivney, B.T. Milliken, G.G. Holz, S.V. Applebey, M.R. Hayes, I.R. Sweet, C.L. Roth, and R.P. Doyle, A peptide triple agonist of GLP-1, neuropeptide Y1, and neuropeptide Y2 receptors promotes glycemic control and weight loss. Scientific reports 13 (2023) 9554.

[13] B.T. Milliken, C. Elfers, O.G. Chepurny, K.S. Chichura, I.R. Sweet, T. Borner, M.R. Hayes, B.C. De Jonghe, G.G. Holz, C.L. Roth, and R.P. Doyle, Design and Evaluation of Peptide Dual-Agonists of GLP-1 and NPY2 Receptors for Glucoregulation and Weight Loss with Mitigated Nausea and Emesis. Journal of medicinal chemistry 64 (2021) 1127–1138.

[14] L.L. Baggio, Q. Huang, T.J. Brown, and D.J. Drucker, Oxyntomodulin and glucagon-like peptide-1 differentially regulate murine food intake and energy expenditure. Gastroenterology 127 (2004) 546–58.

[15] J.E. Blevins, B.W. Thompson, V.T. Anekonda, J.M. Ho, J.L. Graham, Z.S. Roberts, B.H. Hwang, K. Ogimoto, T.H. Wolden-hanson, J.O. Nelson, K.J. Kaiyala, P.J. Havel, K.L. Bales, G.J. Morton, M.W. Schwartz, and D.G. Baskin, Chronic CNS oxytocin signaling preferentially induces fat loss in high fat diet-fed rats by enhancing satiety responses and increasing lipid utilization. Am J Physiol-Reg I (2016).

[16] Z.S. Roberts, T.H. Wolden-Hanson, M.E. Matsen, V. Ryu, C.H. Vaughan, J.L. Graham, P.J. Havel, D.W. Chukri, M.W. Schwartz, G.J. Morton, and J.E. Blevins, Chronic Hindbrain Administration of Oxytocin is Sufficient to Elicit Weight Loss in Diet-Induced Obese Rats. Am J Physiol Regul Integr Comp Physiol (2017) ajpregu 00169 2017.

[17] A.A. Bremer, K.L. Stanhope, J.L. Graham, B.P. Cummings, W. Wang, B.R. Saville, and P.J. Havel, Fructose-fed rhesus monkeys: a nonhuman primate model of insulin resistance, metabolic syndrome, and type 2 diabetes. Clinical and translational science 4 (2011) 243–52.

[18] M.M. Edwards, H.K. Nguyen, A.J. Herbertson, A.D. Dodson, T. Wietecha, T. Wolden-Hanson, J.L. Graham, K.D. O’Brien, P.J. Havel, and J.E. Blevins, Chronic Hindbrain Administration of Oxytocin Elicits Weight Loss in Male Diet-Induced Obese Mice. Am J Physiol Regul Integr Comp Physiol (2021).

[19] K.J. Livak, and T.D. Schmittgen, Analysis of relative gene expression data using real-time quantitative PCR and the 2(-Delta Delta C(T)) Method. Methods 25 (2001) 402–8.

[20] P. Naveilhan, H. Hassani, J.M. Canals, A.J. Ekstrand, A. Larefalk, V. Chhajlani, E. Arenas, K. Gedda, L. Svensson, P. Thoren, and P. Ernfors, Normal feeding behavior, body weight and leptin response require the neuropeptide Y Y2 receptor. Nature medicine 5 (1999) 1188–93.

[21] N.M. Neary, C.J. Small, M.R. Druce, A.J. Park, S.M. Ellis, N.M. Semjonous, C.L. Dakin, K. Filipsson, F. Wang, A.S. Kent, G.S. Frost, M.A. Ghatei, and S.R. Bloom, Peptide YY3-36 and glucagon-like peptide-17-36 inhibit food intake additively. Endocrinology 146 (2005) 5120–7.

[22] B.B. Boland, R.C. Laker, S. O’Brien, S. Sitaula, I. Sermadiras, J.C. Nielsen, P. Barkholt, U. Roostalu, J. Hecksher-Sorensen, S.R. Sejthen, D.D. Thorbek, A. Suckow, N. Burmeister, S. Oldham, S. Will, V.G. Howard, B.M. Gill, P. Newton, J. Naylor, D.C. Hornigold, J. Austin, L. Lantier, O.P. McGuinness, J.L. Trevaskis, J.S. Grimsby, and C.J. Rhodes, Peptide-YY(3-36)/glucagon-like peptide-1 combination treatment of obese diabetic mice improves insulin sensitivity associated with recovered pancreatic beta-cell function and synergistic activation of discrete hypothalamic and brainstem neuronal circuitries. Molecular metabolism 55 (2022) 101392.

[23] K. Tatarkiewicz, E.J. Sablan, C.J. Polizzi, C. Villescaz, and D.G. Parkes, Long-term metabolic benefits of exenatide in mice are mediated solely via the known glucagon-like peptide 1 receptor. Am J Physiol Regul Integr Comp Physiol 306 (2014) R490–8.

[24] M.R. Hayes, K.P. Skibicka, and H.J. Grill, Caudal brainstem processing is sufficient for behavioral, sympathetic, and parasympathetic responses driven by peripheral and hindbrain glucagon-like-peptide-1 receptor stimulation. Endocrinology 149 (2008) 4059–68.

[25] J.P. Krieger, E.P. Santos da Conceicao, G. Sanchez-Watts, M. Arnold, K.G. Pettersen, M. Mohammed, S. Modica, P. Lossel, S.F. Morrison, C.J. Madden, A.G. Watts, W. Langhans, and S.J. Lee, Glucagon-like peptide-1 regulates brown adipose tissue thermogenesis via the gut-brain axis in rats. Am J Physiol Regul Integr Comp Physiol 315 (2018) R708–R720.

[26] H.J. van Eyk, E.H.M. Paiman, M.B. Bizino, I.J. SL, F. Kleiburg, T.G.W. Boers, E.J. Rappel, J. Burakiewicz, H.E. Kan, J.W.A. Smit, H.J. Lamb, I.M. Jazet, and P.C.N. Rensen, Liraglutide decreases energy expenditure and does not affect the fat fraction of supraclavicular brown adipose tissue in patients with type 2 diabetes. Nutrition, metabolism, and cardiovascular diseases : NMCD 30 (2020) 616–624.

[27] S. Gabery, C.G. Salinas, S.J. Paulsen, J. Ahnfelt-Ronne, T. Alanentalo, A.F. Baquero, S.T. Buckley, E. Farkas, C. Fekete, K.S. Frederiksen, H.C.C. Helms, J.F. Jeppesen, L.M. John, C. Pyke, J. Nohr, T.T. Lu, J. Polex-Wolf, V. Prevot, K. Raun, L. Simonsen, G. Sun, A. Szilvasy-Szabo, H. Willenbrock, A. Secher, L.B. Knudsen, and W.F.J. Hogendorf, Semaglutide lowers body weight in rodents via distributed neural pathways. Jci Insight 5 (2020).

[28] J. Blundell, G. Finlayson, M. Axelsen, A. Flint, C. Gibbons, T. Kvist, and J.B. Hjerpsted, Effects of once-weekly semaglutide on appetite, energy intake, control of eating, food preference and body weight in subjects with obesity. Diabetes, obesity & metabolism 19 (2017) 1242–1251.

[29] S.J. Lee, G. Sanchez-Watts, J.P. Krieger, A. Pignalosa, P.N. Norell, A. Cortella, K.G. Pettersen, D. Vrdoljak, M.R. Hayes, S.E. Kanoski, W. Langhans, and A.G. Watts, Loss of dorsomedial hypothalamic GLP-1 signaling reduces BAT thermogenesis and increases adiposity. Molecular metabolism 11 (2018) 33–46.

[30] S.H. Lockie, K.M. Heppner, N. Chaudhary, J.R. Chabenne, D.A. Morgan, C. Veyrat-Durebex, G. Ananthakrishnan, F. Rohner-Jeanrenaud, D.J. Drucker, R. DiMarchi, K. Rahmouni, B.J. Oldfield, M.H. Tschop, and D. Perez-Tilve, Direct control of brown adipose tissue thermogenesis by central nervous system glucagon-like peptide-1 receptor signaling. Diabetes 61 (2012) 2753–62.

[31] Q. Wei, L. Li, J.A. Chen, S.H. Wang, and Z.L. Sun, Exendin-4 improves thermogenic capacity by regulating fat metabolism on brown adipose tissue in mice with diet-induced obesity. Ann Clin Lab Sci 45 (2015) 158–65.

[32] F.C.B. Oliveira, E.J. Bauer, C.M. Ribeiro, S.A. Pereira, B.T.S. Beserra, S.M. Wajner, A.L. Maia, F.A.R. Neves, M.S. Coelho, and A.A. Amato, Liraglutide Activates Type 2 Deiodinase and Enhances beta3-Adrenergic-Induced Thermogenesis in Mouse Adipose Tissue. Frontiers in endocrinology 12 (2021) 803363.

[33] C.M. Mack, C.X. Moore, C.M. Jodka, S. Bhavsar, J.K. Wilson, J.A. Hoyt, J.L. Roan, C. Vu, K.D. Laugero, D.G. Parkes, and A.A. Young, Antiobesity action of peripheral exenatide (exendin-4) in rodents: effects on food intake, body weight, metabolic status and side-effect measures. Int J Obes (Lond) 30 (2006) 1332–40.

[34] K. Erreger, A.R. Davis, A.M. Poe, N.H. Greig, G.D. Stanwood, and A. Galli, Exendin-4 decreases amphetamine-induced locomotor activity. Physiol Behav 106 (2012) 574–8.

[35] C.M. Kotz, C.E. Perez-Leighton, J.A. Teske, and C.J. Billington, Spontaneous Physical Activity Defends Against Obesity. Current obesity reports 6 (2017) 362–370.

[36] M. Periasamy, J.L. Herrera, and F.C.G. Reis, Skeletal Muscle Thermogenesis and Its Role in Whole Body Energy Metabolism. Diabetes Metab J 41 (2017) 327–336.

[37] Q. Yang, W. Tang, L. Sun, Z. Yan, C. Tang, Y. Yuan, H. Zhou, F. Zhou, S. Zhou, Q. Wu, P. Song, T. Fang, R. Xu, J. Han, and N. Jiang, Design of Xenopus GLP-1-Based Long-Acting Dual GLP-1/Y(2) Receptor Agonists. Journal of medicinal chemistry 65 (2022) 14201–14220.

[38] V. Parthsarathy, C. Hogg, P.R. Flatt, and F.P.M. O’Harte, Beneficial long-term antidiabetic actions of N- and C-terminally modified analogues of apelin-13 in diet-induced obese diabetic mice. Diabetes, obesity & metabolism 20 (2018) 319–327.

[39] P.L. Alves, F.M.F. Abdalla, R.F. Alponti, and P.F. Silveira, Anti-obesogenic and hypolipidemic effects of a glucagon-like peptide-1 receptor agonist derived from the saliva of the Gila monster. Toxicon 135 (2017) 1–11.

[40] M.E. Heath, Neuropeptide Y and Y1-receptor agonists increase blood flow through arteriovenous anastomoses in rat tail. J Appl Physiol (1985) 85 (1998) 301–9.

[41] J.P. Frias, M.J. Davies, J. Rosenstock, F.C. Perez Manghi, L. Fernandez Lando, B.K. Bergman, B. Liu, X. Cui, K. Brown, and S.-. Investigators, Tirzepatide versus Semaglutide Once Weekly in Patients with Type 2 Diabetes. The New England journal of medicine 385 (2021) 503–515.

[42] T.D. Muller, M. Bluher, M.H. Tschop, and R.D. DiMarchi, Anti-obesity drug discovery: advances and challenges. Nature reviews. Drug discovery (2021).

[43] O.G. Chepurny, R.L. Bonaccorso, C.A. Leech, T. Wollert, G.M. Langford, F. Schwede, C.L. Roth, R.P. Doyle, and G.G. Holz, Chimeric peptide EP45 as a dual agonist at GLP-1 and NPY2R receptors. Scientific reports 8 (2018).

[44] J.P. Frias, E.J. Bastyr, L. Vignati, M.H. Tschop, C. Schmitt, K. Owen, R.H. Christensen, and R.D. DiMarchi, The Sustained Effects of a Dual GIP/GLP-1 Receptor Agonist, NNC0090-2746, in Patients with Type 2 Diabetes. Cell Metabolism 26 (2017) 343-+.

